# Deep learning reveals functional archetypes in the adult human gut microbiome that underlie interindividual variability and confound disease signals

**DOI:** 10.1101/2025.01.29.635381

**Authors:** Mohamed Meawad, Dalwinder Singh, Alice Deng, Rohan Sonthalia, Evelyn Cai, Vanessa Dumeaux

## Abstract

Understanding the functional diversity of the gut microbiome is essential for decoding its roles in health and disease. Using a deep-learning framework, we identified three functional archetypes defining healthy adult gut microbiomes, each characterized by specific metabolic potentials: sugar metabolism with branched-chain amino acid and cell wall synthesis (Archetype 1), fatty acid and TCA cycle metabolism (Archetype 2), and amino acid and nitrogen metabolism (Archetype 3). Archetype proximity is linked to stability, with Archetype 2 representing the most resilient state, likely due to its metabolic flexibility. Functional diversity emerged as a confounder in disease-associated microbial signatures. In inflammatory bowel disease, we observed archetype-specific shifts, including increased carbohydrate metabolism in Archetype 1-dominant samples and inflammatory pathways in Archetype 3-dominant samples, suggesting distinct opportunities for microbiome-targeted interventions. This framework addresses key challenges in microbiome research, including inter-individual variability and confounding, while providing robust insights into disease-associated functional shifts and microbial ecosystem dynamics.

**Highlights:**

- Adult gut microbiomes are defined by three functional archetypes
- Archetypes reveal distinct metabolic potentials and inform on microbiome stability
- Archetype-specific functional profiles confound disease associations and reveal therapeutic targets
- A deep-learning framework enables robust characterization of microbial functional ecosystems

## Introduction

Host-adapted microbial communities and their collective genomes, termed microbiomes, are shaped by host anatomy and physiology, inhabiting diverse ecological niches across the body.^1–3^ Within these niches, microbiomes engage in dense and dynamic interactions - ranging from mutualism to competition - which critically influence their composition, functionality, stability, and resilience.^4–9^

Microbiomes exhibit variability at multiple scales, including differences between host species, body sites, and individuals. This variability poses significant challenges for experimental design and clinical interpretation, hindering reproducibility and limiting biological discovery.^10^ Furthermore, identifying the magnitude and patterns of these variations is essential for understanding microbiome dynamics and their functional consequences.

In the human gut, the microbiome undergoes dramatic changes from infancy to adulthood, ultimately assembling into a relatively stable ‘climax community’.^11,12^ Despite broad interindividual variation, healthy gut microbiota have been broadly categorized into three ‘enterotypes’ dominated by *Bacteroides*, *Prevotella* or *Ruminococcus*.^13–16^ These configurations are associated with host factors such as diet, health, and metabolic characteristics.^17,18^ Similar patterns, which we term Microbial Configurations (MCs), have been identified in other body sites and across animal species, underscoring the generality of this phenomena.^1,19–22^

Early efforts to identify MCs relied on simple clustering techniques, but these methods often oversimplify the complexity of microbial ecosystems. Continuous or ‘quasi-discrete’ gradients have since been proposed as more accurate representations of microbial variation.^17,23–26^ While the recognition of quasi-discrete compositional MCs and their associated factors represent an important step forward, it raises critical questions about their origins and implications. Are these configurations primarily driven by community composition, or do they reflect functional characteristics that transcend taxonomic variation? Moreover, it remains unclear whether compositional MCs sufficiently capture overall community function or if functional MCs diverge from compositional patterns altogether.

These questions are fundamental to not only understand microbiome inter-individual variability and function but also underlying parameters defining their stability and resilience. Functional redundancy—where multiple species within a community perform overlapping roles—has emerged as a key mechanism underlying microbiome resilience, ensuring the preservation of essential functions despite compositional fluctuations.^27^ Furthermore, metabolic independence and resilience in stressed gut environments have been identified as crucial factors for maintaining host health.^28^ These insights highlight the need for analytical frameworks that can capture the functional dynamics of microbial ecosystems.

To address these questions, we used an analytical framework that combines deep learning and archetypal analysis to model non-linear functional interactions within microbial ecosystems.^29–32^ By identifying extremal points, or archetypes, this approach represents each microbiome’s functional profile as a mixture of archetypes, enabling a nuanced characterization of functional MCs. Using this framework, we define the functional variability of global healthy adult gut microbiomes and investigate the interplay between composition, function, and stability. This approach addresses key challenges in microbiome research, including inter-individual variability and confounding, providing robust insights into disease-associated functional shifts. Beyond the gut microbiome, this framework establishes a foundation for studying microbial ecosystems in diverse contexts, with broad implications for understanding microbial community dynamics and developing microbiome-based interventions.

## Results

### A comprehensive compendium of healthy adult gut microbiomes for robust functional archetype identification

We developed a comprehensive compendium to integrate metagenomic profiles from various large dataset repositories. Key sources include the curatedMetagenomicData R package^33^, GMrepo^34^, and additional controlled-access datasets such as LifeLines DEEP^35^ and Milieu Intérieur^36^ (**Table 1**; **Table S1**). Each sample underwent processing with a computational pipeline designed to use the latest genome annotations and minimize false positives, converting raw reads into relative pathway abundances (**Figure S1; STAR Methods**). We acknowledge that much of the mapping to functional databases likely derives from common housekeeping and well-characterized genes, prevalent across diverse bacterial species and well-represented in reference databases.

**Table 1.** Geographic distribution of curated human gut microbiome datasets. Summary of curated healthy adult gut microbiome datasets after quality filtering based on library size and cohort size. The table shows the number and percentage of profiles from the Global North and Global South regions for each data source.

| Data source | Global North<br>*n* (%) | Global South<br>*n* (%) |
| --- | --- | --- |
| CuratedMetagenomicData<sup>a</sup> | 5319 (82) | 1137 (18) |
| GMrepo V2<sup>a</sup> | 603 (79) | 162 (21) |
| EGA (controlled-access) |  |  |
| Lifelines-DEEP <sup>36</sup> | 1108 (100) | 0 (0%) |
| Milieu Interieur <sup>35</sup> | 1351 (100) | 0 (0%) |
| PubMed |  |  |
| PRJEB49206 (Carter et al., 2023) | 0 (0) | 158 (100) |
| Total | 8381(85%) | 1457 (15%) |

<sup>a</sup> Individual BioProject IDs are provided in Table S1

Our compendium represents the largest repository of whole-genome metagenomic (WGM) profiles from healthy adults, comprising 11,309 samples from 76 studies conducted in 31 countries, mostly from the Global North (**Figure 1A**, **Table 1; STAR Methods**). On average, WGM studies achieved approximately 900,000 reads per sample, with a range from 55,274 to 12,437,351 reads, spanning an average of 334 pathways (range: 1-575) (**Figure 1B-C**). Illumina sequencing was predominantly used in 98.6% of studies, while a small fraction (1.4%) used Ion proton sequencing.

**Figure 1.**
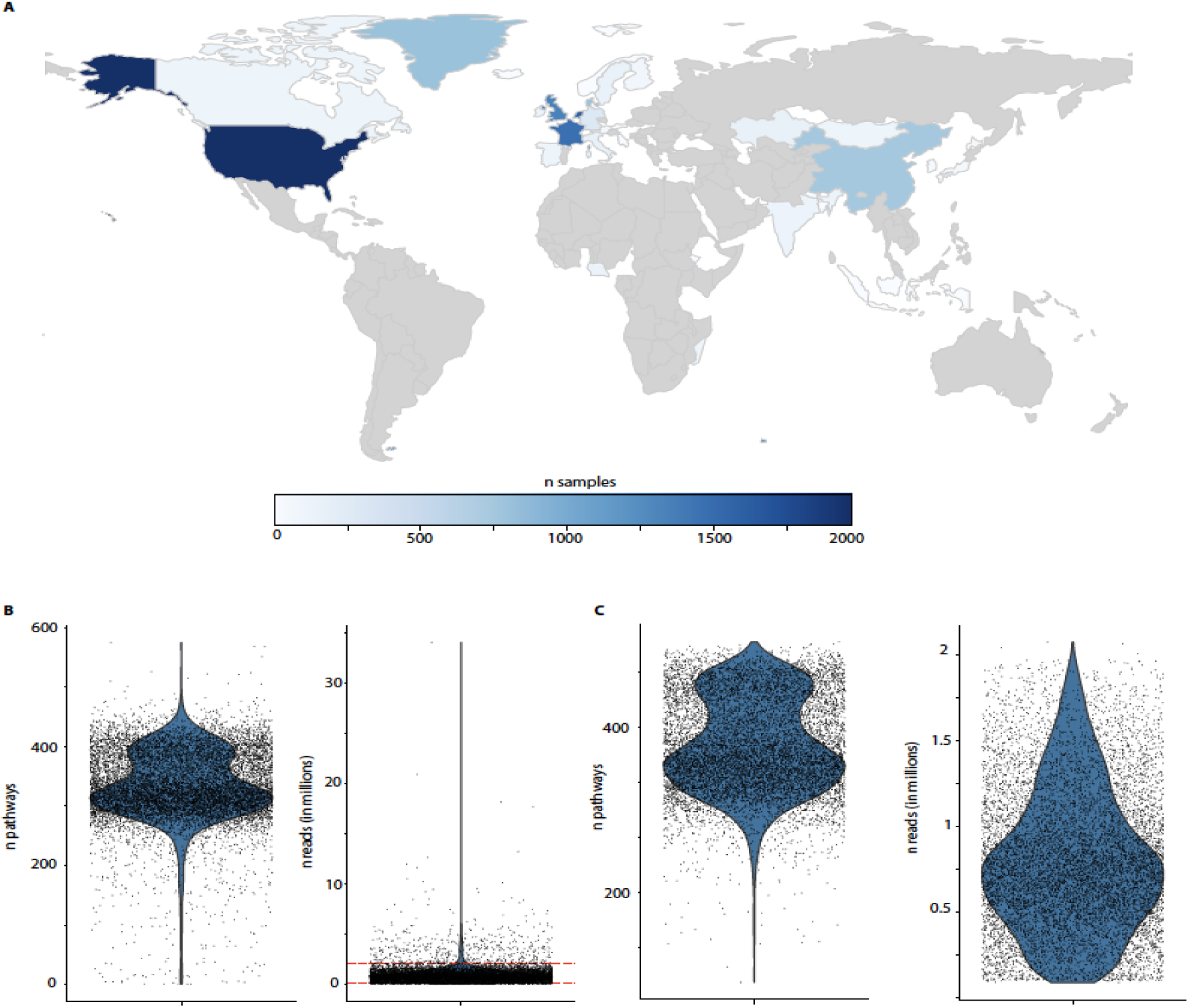
A large compendium of healthy adult microbiomes for robust functional archetype identification. **(A)** A world map illustrating the distribution of microbiome samples across countries before filtering, with the number of samples per country indicated by color. Countries without available samples are shown in gray. **(B-C)** Violin plots displaying the distribution of the number of functional pathways (left) and library size (right) for **(B)** the raw functional pathway data and **(C)** the filtered dataset. Filtering criteria included retaining samples with a total read count (library size) between 100,000 and 2,000,000 reads, removing pathways present in fewer than 10% of samples, and excluding studies with fewer than 30 samples.

To ensure data quality, we excluded samples with total read counts (library size) below 100,000 or above 2 million, as well as studies with fewer than 30 samples. Additionally, pathways with zero counts in more than 10% of samples were removed (**Figure 1B-C)**. The final dataset comprises 436 pathways, with samples expressing an average of 333 pathways (range: 118-427), across 9,838 samples available for downstream analyses (**Figure S2A-B**).

A significant batch effect associated with study origin was detected and corrected using ComBat-Seq^37^ (**Figure S2C-D; STAR Methods**). Notably, no substantial batch effect was observed for Global Region beyond study of origin (**Figure S2E-F**), suggesting that despite the compendium including only 15% of samples from the Global South, the results after adjustment for study are likely generalizable to these populations. Importantly, the batch correction preserved key dataset properties, including the prevalence of zeros and library size of samples in the dataset (**Figure S3**; **STAR Methods**), as well as the count distributions, with the negative binomial (NB) distribution identified as the best fit for our functional relative abundance data (**Figure S4**; **STAR Methods**).

Following batch correction, we applied non-linear archetypal analysis by integrating archetypal analysis with a deep autoencoder^32^ (**Figure S1; STAR Methods**). This approach decomposes each microbiome functional profile into a mixture of archetypes, representing extreme profiles. Model optimization was achieved by minimizing a combined loss function, incorporating the archetype loss (measuring the distance between predicted and fixed archetypes) and the reconstruction loss (using negative log-likelihood NB loss to account for count distribution characteristics) (**STAR Methods**).

Using stability metrics that assess both archetype robustness and sample assignment consistency, we identified K=3 as the optimal number of archetypes **(Figure S5; STAR Methods)**. To ensure reproducibility, we ran the model across multiple random states (n=100) and employed a rigorous selection process based on cosine similarity and archetype distinctiveness to identify the most representative state **(Figure S6; STAR Methods)**. The high stability values and consistent archetype patterns across random states demonstrate the robustness of our three-archetype solution.

To visualize the relationship between samples and their archetypal compositions, we projected the data using multidimensional scaling (MDS), where the three archetypes define the vertices of the solution space. The MDS visualization revealed that while some samples closely aligned with pure archetypes (vertices), the majority of samples showed varying degrees of contribution from multiple archetypes, suggesting that most human gut microbiomes represent functional combinations rather than discrete states **(Figure 2A**). There is a notable density gradient in the archetype usage space, with an enrichment of samples with mid to high usage for both archetypes 1 and 3 (**Figure 2B**). Interestingly, samples with greater affinity to Archetype 2 contained a higher number of detected pathways but it was not reflected in the compositional data providing the number of species identified in the samples (**Figure 2C,D**). We observed differences in the distribution of Archetype 2 scores across age groups and in the distributions of Archetypes 1 and 3 across sexes (ANOVA or t-test, p < 0.001; **Figure S7A**). However, these differences may be influenced by variations in country or regional distributions (**Figure S7B)**, as sex and age are not equally represented across countries.

**Figure 2.**
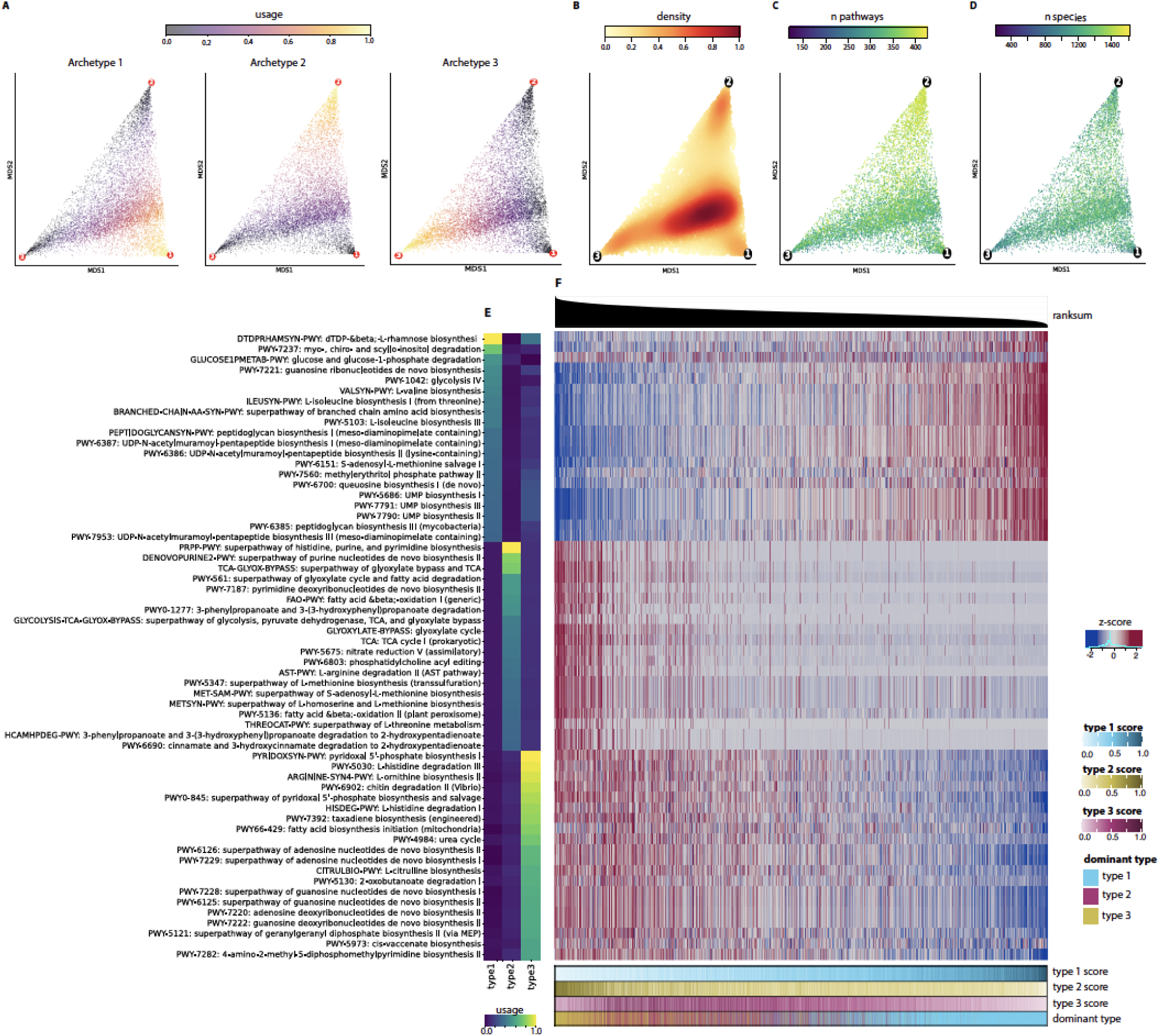
Pathway usage and abundance patterns across archetypal states. **(A)** Multidimensional scaling (MDS) representation of samples based on their archetypal compositions. Each panel shows the same MDS coordinates, with color intensity indicating the relative contribution (usage) of each archetype (scale 0-1). The three archetypes (labeled 1-3) define the vertices of the solution space, and each sample’s position reflects its mixture of archetypal contributions, which sum to 1. Left, middle, and right panels highlight the usage of Archetype 1, 2, and 3, respectively. (**B-D)** Same MDS coordinates as in A), with color intensity indicating **(B**) the sample density or the number of unique (**C)** functional pathways and (**D)** species present in each sample. **(E)** Heatmap showing the relative contribution (usage) of the top 20 pathways in defining each archetype identified by the deepAA model. Color intensity indicates the degree of pathway usage (scale 0-1) for each archetype (type 1-3). (**F)** Heatmap displays pathway abundances across samples for the top 20 pathways defining each archetype. Values were scaled and winsorized at the 2nd and 98th percentiles (z-score). Samples are ordered by relative pathway abundance average ranksum plotted on top of the heatmap. Bottom annotations show archetypal scores and dominant archetype classification for each sample, where dominant type is determined by the highest score among the three archetypes.

### Gut microbiome functional states represent blends of three distinct metabolic archetypes

To define the functional characteristics of each archetype, we identified the top 20 pathways that most strongly distinguished them (**Figure 2E, Table S2**). Each archetype displayed distinct metabolic signatures, reflecting their unique functional roles. To explore how these signature pathways manifest across samples, we visualized their abundance patterns in relation to archetypal classifications (**Figure 2F**). The resulting heatmap revealed gradual transitions in pathway abundances across samples, aligning with their archetypal scores.

Archetype 1 is characterized by high potential for carbohydrate sugar metabolism and biosynthesis of branched-chain amino acid (BCAA) and microbial cell wall component biosynthesis (**Figure 2E, Table S2**). The dTDP-β-L-rhamnose biosynthesis pathway showed maximal enrichment, accompanied by other carbohydrate-processing pathways, including glycolysis IV, glucose and glucose-1-phosphate degradation, and myo-, chiro-, and scyllo-inositol degradation, which collectively constituted four of the top five pathways defining this archetype (**Figure 2E, Table S2**). These sugar metabolism pathways supply key intermediates—such as D-glyceraldehyde 3-phosphate, pyruvate, and phosphoenolpyruvate—that can fuel multiple downstream processes also highly represented in this archetype (**Figure 3A**). For instance, pyruvate serves as a critical substrate for highly expressed BCAA biosynthesis pathways and methylerythritol phosphate pathway. Similarly, phosphoenolpyruvate contributes to peptidoglycan biosynthesis pathways essential for microbial cell wall. In addition to sugars, this archetype relies on L-glutamate as a key substrate for several highly expressed pathways involved in BCAA biosynthesis and peptidoglycan production (**Figure 3A**). L-glutamate is supplied by pathways such as S-adenosyl-L-methionine salvage I, guanosine ribonucleotides de novo biosynthesis, and UMP biosynthesis pathways. Together, these metabolic features highlight Archetype 1’s specialized role in carbohydrate utilization and biosynthesis of key structural and functional components, including BCAAs and components of microbial cell wall such as peptidoglycans.

**Figure 3.**
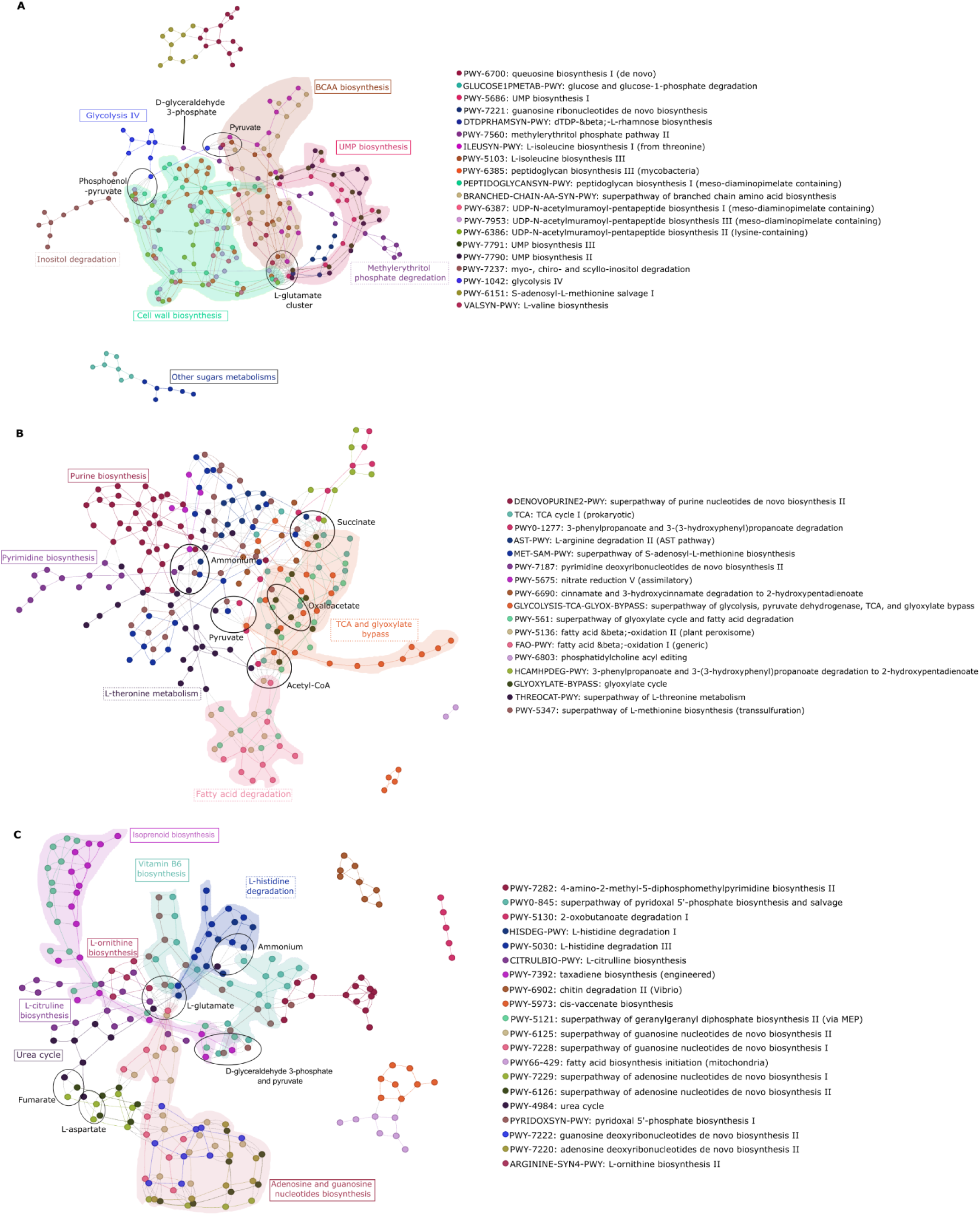
Pathways defining each archetype focuses on specific metabolic strategies favoring different cellular functions. Graphs of the compounds involved in the reactions of the top 20 pathways characterizing **(A)** Archetype 1, **(B)** Archetype 2 and **(C)** Archetype 3 (**STAR Methods**). Each pathway and its nodes are given a single color that matches the legend on the right. Each node represents a single compound in a pathway’s reactions. Nodes / compounds that are duplicates, i.e. found in multiple pathways, are connected with a dashed black line to help visualize core compounds in the archetype and similarities between the pathways’ reactions. Label boxes were added to highlight the core components defining each archetype. Common compounds, such as H+, phosphate, ATP, ADP, H2O, NADP+, NADPH, NADH, NAD+, CO2, coenzyme A, AMP, dioxygen, hydrogen carbonate, and diphosphate, were excluded to reduce visual noise. Interactive graph html files can be found in https://osf.io/tvu52/

Archetype 2 is defined by pathways integrating glycolysis, TCA cycle, glyoxylate bypass, and fatty acid metabolism, driving the production of acetyl-CoA, succinate, oxaloacetate, and pyruvate (**Figure 2E**, **Figure 3B, Table S2**). Central carbon metabolism integrates glycolysis, the TCA cycle, and related pathways to process carbon substrates into energy (ATP) and biosynthetic precursors such as acetyl-CoA and oxaloacetate. Archetype 2’s elevated potential activity within central carbon metabolism, including pyruvate dehydrogenase, underscores the integration of multiple pathways to meet both energy and biosynthetic demands. Acetyl-CoA is both produced and consumed across diverse pathways, including fatty acid degradation, L-threonine metabolism, and aromatic compound degradation, while its consumption in the glyoxylate bypass generates succinate as a key downstream product (**Figure 3B**). Succinate production is a recurring feature of Archetype 2, supported by amino acid degradation, methionine biosynthesis, and central carbon metabolism, with some pathways demonstrating a metabolite-dependent ability to both produce and consume succinate, while others, such as nucleotide biosynthesis, exclusively consume it (**Figure 3B**). Finally, ammonium is produced through pathways linked to amino acid and nucleotide metabolism, while oxaloacetate and pyruvate can be both produced and consumed depending on availability (**Figure 3B**). Unlike Archetype 1, which supports branched-chain amino acid biosynthesis, Archetype 2 prioritizes energy metabolism and nucleotide biosynthesis, emphasizing its role in driving core biosynthetic and energy-yielding processes.

Finally, the top 20 pathways representing Archetype 3 reveal links between the urea cycle (including the biosynthesis and recycling of intermediates like L-ornithine and L-citrulline), nucleoside biosynthesis, isoprenoid biosynthesis, and vitamin B6 biosynthesis (**Figure 2E**, **Figure 3C, Table S2**). Specifically, the urea cycle and histidine degradation pathways are indirectly connected to nucleoside biosynthesis through key intermediates such as fumarate/L-aspartate and L-glutamate, respectively, which integrate nitrogen and carbon metabolism (**Figure 3C**). Histidine degradation produces ammonium, which is further processed in the urea cycle, creating a direct link between these pathways in nitrogen metabolism. Isoprenoid biosynthesis and vitamin B6 pathways share the same initial compounds such as D-glyceraldehyde 3-phosphate and pyruvate (**Figure 3C**). Vitamin B6 is a cofactor essential for numerous enzymatic processes, including those in the urea cycle, amino acid metabolism and nitrogen metabolism (**Figure 3C**). Overall, this network demonstrates the tight metabolic integration between nitrogen metabolism through the urea cycle and histidine degradation and the production of essential components and cofactors (nucleosides and vitamin B6).

Together, these archetypes highlight specific metabolic strategies which might favor different ecosystems and functions with different impacts on the host’s physiology and health: Archetype 1 may favor microbes that thrive on abundant carbohydrates supporting structural and essential cellular component biosynthesis for microbial growth (microbial cell wall, BCAA). Archetype 2 may favor microbes that use multiple energy harvesting routes supporting metabolic flexibility with specialization in fatty-acid metabolism. Finally, Archetype 3 may favor microbes with enhanced nitrogen metabolism capabilities, particularly in processing nitrogen compounds (including amino-acids) through urea cycle and related pathways.

### Functional archetypes display distinct associations with gut microbiome compositional enterotypes

To investigate potential relationships between functional archetypes and compositional community structures, we analyzed enterotype distributions across our cohort and functional archetypes. Using the recent Enterotyper tool^24^, we first classified samples into three enterotypes defined by predominant taxa: *Bacteroides/Phocaeicola*, *Prevotella*, or *Firmicutes* (**STAR Methods**). While the Firmicutes enterotype was the most prevalent in our dataset (55 %, 32% Bacteroides, 12% Prevotella), samples classified as Prevotella enterotype showed notably stronger classification scores (median classification strength 0.69, 0.55 for Bacteroides, 0.54 for Firmicutes), suggesting more distinct compositional profiles (**Figure 4A**).

**Figure 4.**
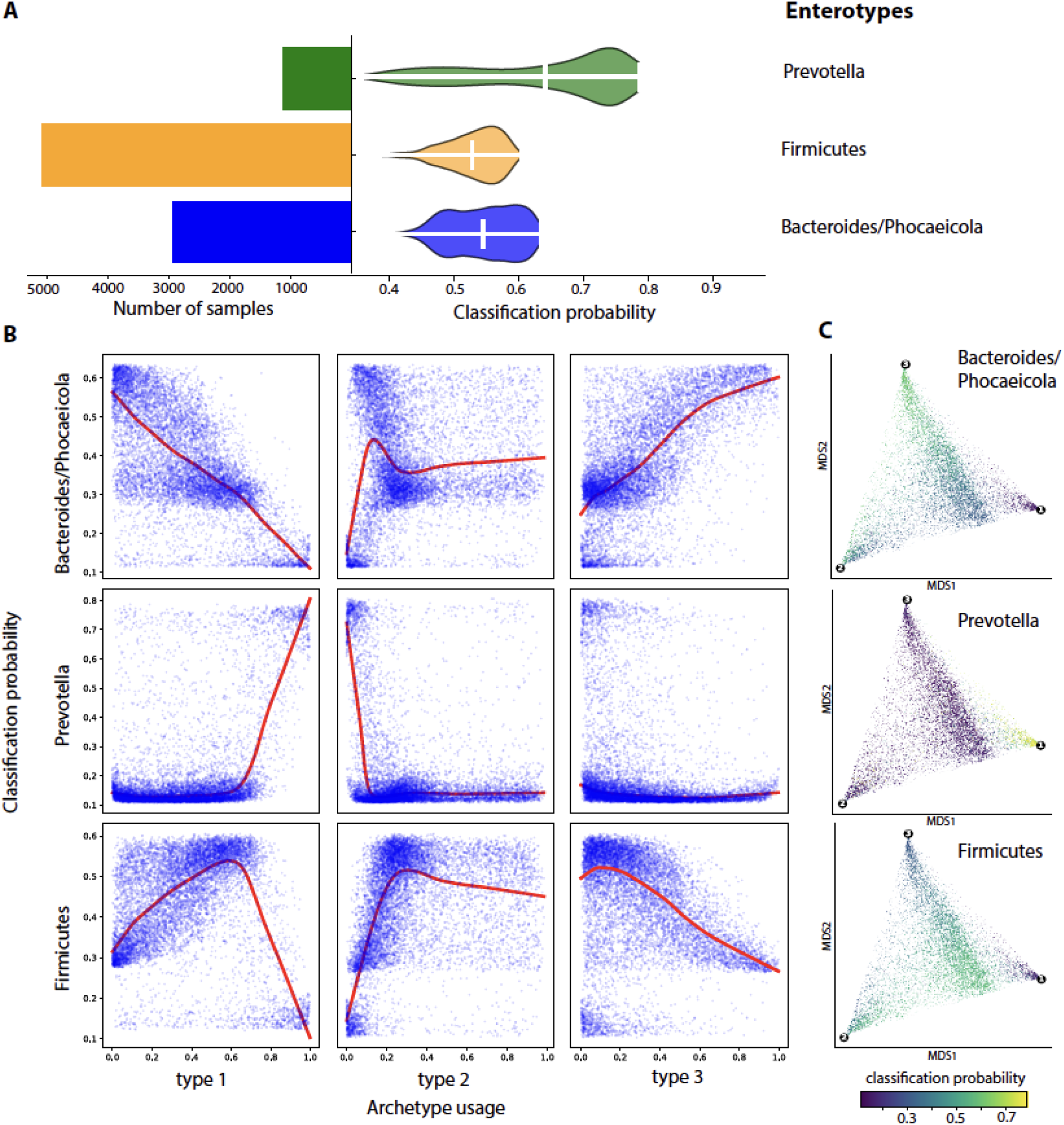
Functional archetypes display distinct associations with gut microbiome compositional enterotypes. **(A)** Number of samples and classification probability for each enterotype **(B)** Scatterrplots depicting relationships between usage of each archetype and classification probability of each enterotype. Lowess curves are depicted in red. **(C)** Multidimensional scaling (MDS) representation of samples based on their archetypal compositions colored by the classification probability of each enterotype.

Our analysis revealed significant but not exclusive associations between functional archetypes and compositional enterotypes. Notably, samples exhibiting high scores for archetype 1 (characterized by enhanced carbohydrate metabolism and BCAA and cell wall component biosynthesis potential) showed strong correlation with the Prevotella enterotype classification (**Figure 4B-C)**. A similar but less exclusive pattern emerged between archetype 3 (elevated nitrogen and amino acid metabolism potential) and the Bacteroides/Phocaeicola enterotype (**Figure 4B-C)**. Samples classified with higher probability as Firmicutes enterotype clustered primarily in the densely populated region of the archetypal space, characterized by either elevated archetype 1 scores with intermediate archetype 3 scores, or high archetype 2 scores (distinguished by enhanced energy-yielding potential, particularly in fatty acid metabolism and central carbon pathways).

Overall, these results indicate that while specific associations exist between functional archetypes and enterotypes, the relationship is not strictly deterministic. The observed patterns suggest that similar functional capabilities can be maintained across different compositional configurations in the adult human gut microbiome.

### Functionally diverse archetype 2 shows enhanced temporal stability

To assess the stability of archetypal states, we analyzed the variation in archetype contributions of samples collected across consecutive visits from the same individuals (7 studies; n subjects = 656; n samples = 1557; range visits 2-6 in the span of 2 - 730 days; **Table S3**). Notably, archetype 2, which is characterized by high pathway diversity and enrichment in energy-yielding processes, showed more consistent scores between consecutive samples, with changes distributing more tightly around zero compared to the broader variations seen in archetypes 1 and 3 (**Figure 5A**).

**Figure 5.**
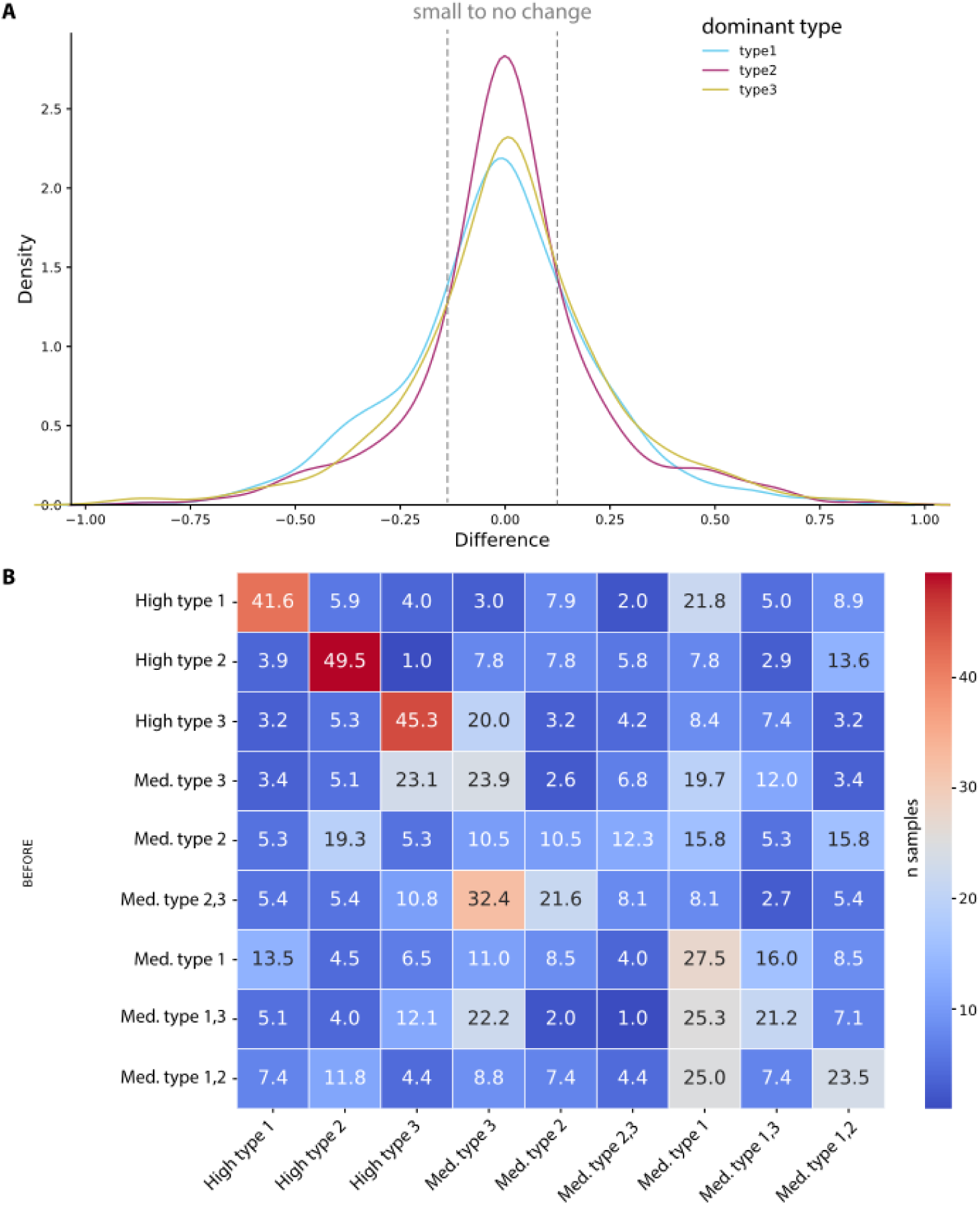
Temporal dynamics of functional archetype scores in the human gut microbiome. **(A)** Density plot showing the difference between samples’ archetype values that are from the same subject. Colors reflect scores for the different archetypes: type 1 (blue), type 2 (red) and type 3 (yellow) **(B)** Heatmap showing the probability of transitioning between archetype states for subjects with multiple samples. Sample states were categorized based on their archetype usage levels: high type usage (≥ 0.66 archetype value), if they did not have a high type usage then they were set to medium type usage (0.33–0.66 archetype value).

We further classified samples into high (≥0.66) and medium (0.33-0.66) archetype usage categories to examine state transitions. Individuals exhibiting high usage of a particular archetype showed substantial persistence of that state, with maintenance percentages of 41.6%, 49.5%, and 45.3% for archetypes 1, 2, and 3, respectively (**Figure 5B**). In contrast, samples with medium usage levels showed lower state stability, with persistence percentages ranging from 8% to 27.5%. These findings suggest that while strongly committed archetypal states tend to persist over time, microbiomes with intermediate archetype contributions display greater temporal flexibility in their metabolic configurations.

### Functional archetypal space captures disease-associated microbiomes and reveal key confounders

To determine whether the archetypal space captures disease-associated gut microbiomes, we curated metagenomic profiles from studies of patients with inflammatory bowel disease (IBD; 3 studies), type 2 diabetes (T2D; 3 studies), and colorectal cancer (CRC; 5 studies) (**Table S4A**; **STAR Methods**). Using the model trained on healthy samples, we mapped the disease-associated samples and their respective controls onto the archetypal space (**Figure 6A**). Notably, these samples aligned within the same archetypal framework, but their distributions differed significantly from those of healthy controls included in the same studies (**Figure 6B**). IBD samples exhibited higher usage of Archetype 2 compared to their controls, while CRC samples showed significantly lower Archetype 2 usage (Kolmogorov–Smirnov test, p ≤ 0.05). Similarly, T2D samples displayed distinct distributions compared to healthy controls, with significant differences in the usage of both Archetype 1 and Archetype 3 (Kolmogorov–Smirnov test, p ≤ 0.05).

**Figure 6.**
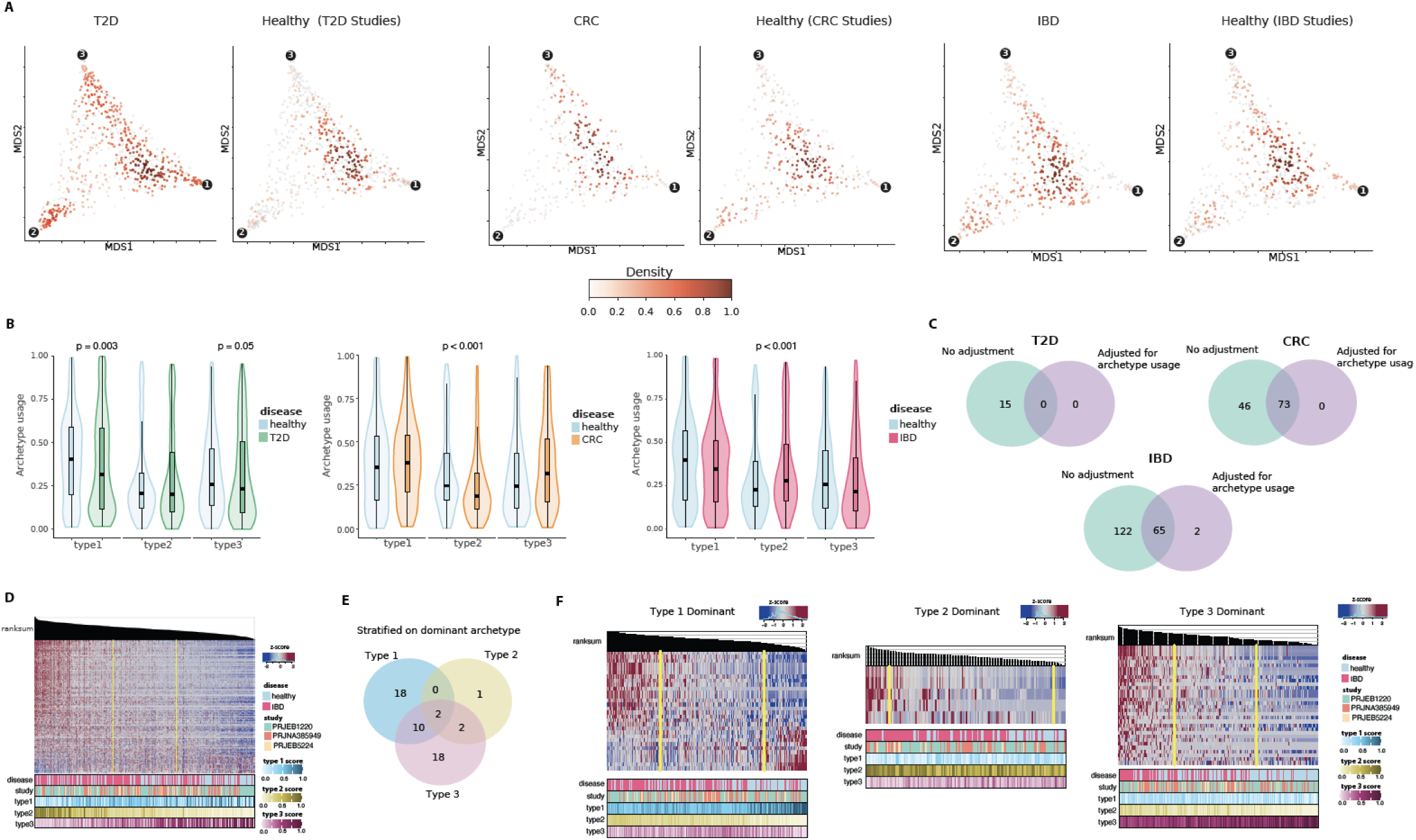
**(A)** Density plots of healthy samples (left) and diseased samples (right) from type 2 diabetes (T2D), colorectal cancer (CRC) and inflammatory bowel disease (IBD) studies, shown in archetypal space after Multidimensional Scaling (MDS). Samples of the opposite health status are displayed in grey. **(B)** Violin plot of archetype usage in diseased study samples, stratified by dominant archetype and colored by disease status. **(C)** Heatmaps of pathways found to be differentially expressed (FDR < 0.01) using MaAslin2, with samples ordered by pathway expression. **(C)** Venn diagrams comparing the number of differentially expressed pathways between healthy and diseased samples from T2D with and without adjustment for archetypes usage. Pathway names are listed in **Table S4B-F. (D)** Heatmaps of pathways differentially expressed (FDR < 0.01, **Table S4E**) between healthy and IBD samples. Samples are ordered based on the ranksum of pathways relative abundance (z-score) depicted on top of heatmap. Metadata at the bottom of the heatmap provide health status and sample’s usage scores for each archetype **(E)** Venn diagram showing the count of differentially expressed pathways (FDR < 0.01) between healthy and IBD samples, stratified by dominant archetype. **(F)** Heatmaps of pathways differentially expressed (FDR < 0.01) between healthy and IBD samples, stratified by dominant archetypes 1 (left), 2 (middle), and 3 (right). Pathway names are listed in **Table S4G-I**.

Given the functional differences between archetypes, we next investigated whether archetype usage could confound the identification of differentially represented pathways between disease and healthy states (**STAR Methods**). Across all diseases (IBD, T2D, and CRC), many pathways identified as differentially represented between disease and healthy samples were also strongly associated with archetype usage. Adjusting for archetype usage significantly reduced the number of differentially enriched pathways (**Figure 6C; Table S4B-F**). For instance, when comparing IBD to healthy samples, pathways initially identified as differentially enriched (n = 187, FDR < 0.01) also largely reflected Archetype 2 usage, further underscoring the confounding influence of archetype-specific functional variability (**Figure 6D**).

Further investigation revealed a significant interaction between IBD status and archetype dominance, defined here as the archetype with highest usage per sample (**Figure 6E-F; Table S4G-I**). Stratified analysis based on archetype dominance revealed distinct disease-associated pathways varying by dominant archetype (**Figure 6E-F; Table S4G-I**).

In IBD samples dominated by Archetype 1 - characterized by high carbohydrate metabolism potential - we identified 32 significant pathways including 18 unique to this archetype (**Figure 6E; Table S4**). Unique pathways included processes related to thiamine biosynthesis, a process critical for carbohydrate metabolism and the production of short-chain fatty acids (SCFAs). Additionally, molybdopterin biosynthesis, essential for anaerobic respiration in bacteria, was significantly overrepresented in the gut microbiome of IBD patients compared to controls. This pathway is particularly relevant given its potential role in the overgrowth of *Enterobacteriaceae* in the inflamed gut.^38^

Conversely, IBD samples dominated by Archetype 3 - associated with high processing capabilities of nitrogen compounds (including amino-acids) - exhibited unique enrichment of pathways involved in NAD salvage, the biosynthesis of amino-acid (L-cysteine, L-glutamine, arginine and polyamines), and fatty-acid related pathways, including one specific to *Escherichia coli*. Fatty-acids play important roles for bacterial membrane synthesis while peptidoglycan biosynthesis essential for bacterial cell wall integrity, was also uniquely overrepresented in gut microbiomes of IBD patients in this archetype (**Figure 6F**).

Notably, ten differential pathways were shared among IBD samples closer to Archetype 1 or 3, including those involved in the TCA cycle, nucleotide degradation, starch metabolism, and heme b biosynthesis. In the unstratified analysis, differential pathways between IBD and controls captured Archetype 2, with IBD samples more likely to have high Archetype 2 usage characterized by high energy-harvesting potential through the TCA cycle. The consistent identification of TCA cycle pathway suggests that dysregulation of central energy metabolism is a common feature across IBD samples, regardless of archetype dominance.

Overall, these findings reveal substantial inter-individual variability in the functional metabolic landscape of gut microbiomes, presenting a significant confounding factor in differential analyses when comparing disease and control groups. The archetypal framework we developed provides a robust approach to mitigating these confounding effects, reducing false positives, and uncovering archetype-specific functional changes. These insights, for example, offer a novel perspective on IBD gut microbiome subtypes and their specific or common metabolic alterations depending on their overall archetypal functional landscape.

## Discussion

Our deep archetypal model revealed that global gut microbiome functional potential can be represented by three archetypes, each defined by a high potential to express pathways within specific metabolic frameworks. Specifically, the three archetypes are skewed toward distinct metabolic features: sugar-related metabolism, whose products feed into branched-chain amino acid (BCAA) and cell wall component biosynthesis (Archetype 1); fatty acid metabolism, whose products fuel the TCA and glyoxylate cycles (Archetype 2); and amino acid metabolism and nitrogen metabolism through the urea cycle and related pathways (Archetype 3).

While most gut microbiome communities are a blend of these archetypes, some communities align closely with a single archetype, potentially reflecting adaptation to specific environmental or host-related conditions. For example, a diet consistently rich in complex carbohydrates might promote a community resembling Archetype 1, while a diet rich in fatty acid or protein might promote a community resembling Archetype 2 and 3, respectively. Similarly, host genetic, physiological factors or disease could create conditions that favor one archetype over others. In some cases, a community dominated by a single archetype could represent a functionally specialized microbiome optimized to meet the host’s needs.

Our findings reveal a nuanced relationship between functional archetypes and compositional enterotypes in the human gut microbiome. While specific associations exist between functional archetypes and enterotypes, the relationship is not strictly deterministic. The strong association between Prevotella enterotype and Archetype 1 reflects biological patterns consistent with the literature, as *Prevotella*-dominant communities are known for their enhanced capacity to metabolize plant-derived carbohydrates and produce SCFAs from diets rich in complex carbohydrates and dietary fibers.^18,39^ Similarly, the connection between Bacteroides/Phocaeicola enterotype and archetype 3’s elevated amino acid metabolism potential corresponds with this enterotype’s known association with protein-rich Western diets.^18,39^ However, compositional configurations still show some flexibility in their functional profiles, supporting the concept of functional redundancy in microbial ecosystems, where different community structures can achieve similar metabolic capabilities. These insights have important implications for microbiome-based interventions, suggesting that targeting functional capabilities rather than specific taxonomic compositions might be a more robust approach for therapeutic strategies.

Proximity to archetypal states confers functional stability of gut microbial communities. In particular, Archetype 2 represents the most stable and diverse state, likely due to its diverse energy harvesting capabilities including fatty acid metabolism, the TCA cycle, and the glyoxylate shunt. This metabolic flexibility through multiple energy-generating pathways could provide enhanced resilience to dietary fluctuations and day-to-day environmental perturbations.

The relationship between archetypal extremes and ecosystem stability presents intriguing questions about the functional stability of disease states. While disease samples are represented within the same archetypal space, they do not necessarily occupy extreme positions nor reflect simple imbalances in archetype usage. Instead, we observed disease-specific enrichments, such as IBD’s association with Archetype 2. This finding aligns with previous findings linking disruptions in the TCA cycle and its intermediates, such as succinate, to heightened inflammation and IBD pathogenesis.^40,41^ However, these patterns warrant careful interpretation, as they might reflect either true functional shifts favoring disease states or sampling biases due to the relatively limited scale and diversity of existing case-control studies. Such biases could also stem from underrepresentation of diseases across the full archetypal space.

Importantly, across all curated disease studies, the distributions of disease samples within the archetypal space consistently differed significantly from those of healthy groups. These differences, rooted in the distinct metabolic profiles of the archetypes, can introduce confounding when directly comparing disease and healthy samples. Our findings further revealed interactions between IBD-associated changes and archetypal usage, consistent with previous findings identifying subtype-specific gut microbiome signatures in IBD patients.^42–45^ These signatures may reflect differences in clinical presentation (e.g., constipated vs non-constipated) or disease stages (e.g., quiescent vs inflamed). For instance, Gargari et al. identified a subgroup of non-constipated IBD patients with higher levels of SCFAs which aligns with our findings in Archetype 1-dominant IBD samples. Archetype 1 is characterized by high carbohydrate metabolism potential and these IBD samples exhibited enrichment for thiamin biosynthesis, a pathway critical for SCFA production through the bacterial fermentation of carbohydrates. These archetype-specific functional changes could inform microbiome-targeted interventions, such as dietary strategies aimed at lowering the fiber-fermenting microbial components of the gut microbiome. Notably, low-fiber diets have previously been shown to be potentially more effective in IBD patients with high fecal SCFA levels.^43,45^ Similarly, our findings highlight the potential for targeting specific metabolic pathways in Archetype 1-dominant IBD samples. For example, tungstate has been shown to prevent *Enterobacteriaceae* overgrowth in the inflamed gut by replacing molybdenum in the molybdopterin cofactor.^38^ This substitution disrupts molybdopterin-dependent enzymatic pathways, which are essential for anaerobic respiration in *Enterobacteriaceae*—a pathway we identified as uniquely enriched in Archetype 1-dominant IBD samples.

In contrast, Archetype 3-dominant IBD samples exhibited enrichment of pathways involved in known inflammatory processes, including immune responses linked to E. coli adherence. These pathways include NAD salvage^46,47^, amino acid biosynthesis^48–50^, and the production of immunogenic cell wall-derived molecules, which may also promote the proliferation of adherent-invasive *Escherichia coli*.^51–53^ These findings underscore the distinct functional and inflammatory mechanisms associated with different archetypes in IBD microbiomes. Overall, our findings underscore the need for large-scale studies with comprehensive sampling of diverse functional microbiomes. Such studies could improve our understanding of the full archetypal landscape and help address potential sampling biases. Incorporating archetype values as a confounding variable or stratification factor in differential analyses reduces inter-individual variability, mitigates confounding, and reveals novel pathways specifically altered in IBD gut microbiomes based on the dominant functional archetypes.

### Limitations

Since our data is derived from whole genome sequencing (WGS) of microbial DNA, the biological archetypes we identified are based on the functional potential encoded within the genomes of the gut microbiome, rather than the actual functional activity occurring at the time of sampling. This means our analysis reflects the possible capabilities of the gut microbiome based on its genetic composition, but it does not capture the dynamic functional state that would be observed through techniques like metatranscriptomics or metaproteomics, which measure RNA transcripts and proteins, respectively.

## Supplemental information

**Document S1.** Figures S1–S3 and Tables S1 – S3

**Table S4.** Excel file containing **(A)** Summary statistics of studies with disease subjects. (B-I) list of differentially expressed pathways between **(B)** healthy and type 2 diabetes samples without archetype values adjustment **(C, D)** healthy and colorectal cancer samples **(C)** without archetype value adjustment **(D)** with archetype value adjustment. **(E, F)** healthy and inflammatory bowel disease samples **(E)** without archetype value adjustment **(F)** with archetype value adjustment. **(G-I)** healthy and inflammatory bowel disease samples when stratifying samples by dominant archetype **(G)** archetype 1 **(H)** archetype 2 *(I)* archetype 3, related to figure 6.

### Lead Contact

Further information and requests for resources should be directed to and will be fulfilled by the lead contact, Vanessa Dumeaux

### Materials availability

This study did not generate new unique reagents.

### Data and code availability

The code used for processing of the raw data is available at https://github.com/dumeaux-lab/compendium-fMC.

The functional profile dataset and the code used for data analyses and generation of the figures in this manuscript is available through the Open Science Foundation (OSF) repository (https://osf.io/tvu52/) and its associated github repository https://github.com/dumeaux-lab/deep-fMC_paper.

## Acknowledgements

This work was supported by Natural Sciences and Engineering Research Council of Canada (NSERC) Discovery grant #391682. We thank the Canadian Foundation for Innovation J. Evans Leaders Fund grant #43481 for the support of the computing infrastructure used throughout our analyses.

MM received the Vector Scholarship in Artificial Intelligence, provided through the Vector Institute and RS received the MITACS Globalink Summer Internship Award.

## Author contributions

Conceptualization, V.D.; methodology, M.M, R.S., V.D.; software, M.M, D.S., V.D.; investigation, M.M., R.S., A.D., V.D.; visualization, M.M., V.D.; funding acquisition, V.D.; data curation, M.M., D.S., E.C., V.D.; supervision, V.D.; writing – original draft, M.M., V.D.; writing – review & editing, M.M., A.D., R.S., E.C., A.D., and V.D.

## STAR Methods

### Compendium data curation, preprocessing, transformation, and filtering

We curated 11309 publicly available whole genome sequencing (WGS) samples’ data from healthy adult (age ≥ 18) stool samples using several large dataset repositories such as GMrepo and curatedMetagenomicData R package as well as datasets under controlled access, namely LifeLines DEEP (EGAR) and Milieu Interieur (WGM) (**Figure 1A**, **Table 1, Table S1**).

Each sample underwent processing with a computational pipeline designed to use the latest genome annotations and minimize false positives, converting raw reads into relative pathway abundances (**Figure S1**). First, the raw reads were preprocessed for quality control and adapter removal using fastp^54^ with the following parameters: `--trim_poly_x --trim_poly_g -p --length_required 40 --cut_front --cut_tail --cut_mean_quality 25`. Based on our experience^55^ and corroborated by previous studies^56,57^, when using properly defined thresholds and a comprehensive reference database, the Kraken2+Bracken^58,59^ toolset delivers superior performance to estimate microbial composition of gut microbiomes. We therefore used Kraken2 with confidence threshold of 0.15 to align reads to the HumGut database^60^ following the Genome Taxonomy Database (GTDB) classification scheme^61^ and the human genome downloaded from NCBI to identify and remove contamination.

Biochemical pathways/functions were then identified using HUMAnN3 v3.7^62^ based on species identified using Kraken2+Bracken and HUMAnN3 reference databases including the UniRef90 and protein GTDB databases obtained from Struo2.^63^ HUMAnN3 quantified pathways in units of RPKs (reads per kilobase) and counts were further normalized for library size where read counts for each sample are constrained to sum to 1 millions (copies per millions, CoPM).

Functional pathways data samples were filtered for total read count (library size) of 100,000 to 2,000,000 reads filter and functional pathways that are present in less than 10% of the samples were removed along with studies with less than 30 samples, 9838 samples were left after filtering (**Figure 1B-C, Figure S2**).

As expected, we observed a significant batch effect associated with sample study source **(Figure S2C)**, and applied batch correction using ComBat Seq^37^ (**Figure S2D**). To further confirm appropriate batch effect adjustment, we compared the assigned archetype values of the second largest study in the dataset (LifeLines DEEP / EGAR, n = 1131), which exhibited the most significant batch effect, to samples from other studies (**Figure S2C**). The violin plot illustrates that there is no bias between the study source of the samples and how each archetype defines a sample, unlike when batch correction is not performed, indicating the effectiveness of ComBat’s batch correction (**Figure S3A-B**). Finally, to ensure that this correction did not alter the distribution or sparsity of the data, a comparison between values in the original and batch-corrected matrices was conducted. Overall, the values in both datasets were overwhelmingly similar, and the sparsity (zero counts) of the data remained identical **(Figure S3C-E)**.

To determine the compositional enterotypes of our samples we used the recent Enterotyper webtool^24^ by submitting the microbial species raw counts and exporting the classification probability table. Microbial species raw counts were filtered for species that were present in at least 10% of samples, filtered against studies with less than 30 samples and batch corrected using ComBat-seq.^37^

### Statistical distribution of pathway abundance

Following batch correction, non-linear archetypal analysis was performed by integrating archetypal analysis with a deep autoencoder (**Figure S1**). Prior to model training, the distribution of the count data was assessed to inform the parameterization of the decoder. Pathway abundance data (RPK) were modeled using the generalized additive model for location, scale, and shape (GAMLSS) as implemented in the scDesign3 R package.^64^ The Akaike Information Criterion (AIC) values were calculated for Poisson, zero-inflated Poisson (ZIP), Gaussian, negative binomial (NB), and zero-inflated negative binomial (ZINB) distributions to evaluate their fit to the data.

NB and ZINB had the lowest AIC values (**Figure S4A-B**), and simulated data using these distributions (**Figure S4C**) produced UMAP visualizations with a structure and distribution qualitatively similar to the original dataset.. Ultimately, we selected NB to parameterize our decoder, as it is less complex, supports Combat-seq batch correction assumptions, and provides a comparable fit to the data than ZINB.

### Deep Archetypal Analysis

We used deep archetypal analysis as implemented in scAAnet.^65^ The total loss function comprised archetypal loss and reconstruction loss. Archetypal loss measured the deviation between fixed archetypes and inferred archetypes in the latent space, while reconstruction loss was calculated as the negative log-likelihood of the NB distribution using model-estimated mean and dispersion parameters.

The scAAnet model was configured with a batch size of 64 and a hidden layer width of 128. The activation function used was a rectified linear unit (ReLU). The model was trained for up to 1000 epochs (each epoch representing one full training cycle), with 20 warm-up epochs. Early stopping was enabled if the loss did not improve for 100 epochs, and the learning rate was reduced if the loss failed to improve for 10 epochs. The initial learning rate was set to 0.01. To identify the optimal number of archetypes (K value) for the data, the model was tested with K values ranging from 2 to 7. Archetype and usage stability were assessed across 8 random states (0 to 7). The dispersion parameter was set to be unique for each pathway.

### Number of archetypes (K value) selection

Two metrics are used to determine the optimal K value (number of archetypes), archetypes’ stability and samples’ archetype usage consistency^65^, as implemented in scAAnet. Archetype stability was assessed by performing K-means clustering on the archetype spectra derived from multiple random initializations (n = 8) and computing the Euclidean distance silhouette score to evaluate the quality and robustness of the inferred archetypes at each K value. This stability metric ranges from 0 to 1, with higher values indicating more robust archetypes.

To assess consistency in sample archetype usage, we identified the dominant archetype for each sample across multiple random states (n = 8). For each pair of samples, we calculated the proportion of times they were assigned to the same archetype, resulting in a consensus matrix C that indicates the probability of two samples being assigned to the same archetype. Using these pairwise probabilities as measures of similarity, we performed average linkage hierarchical clustering to reorder the samples. We then calculated the cophenetic distance for each pair of samples based on the hierarchical clustering dendrogram, reflecting how similar two samples are within the dendrogram. The cophenetic correlation coefficient (ranging from 0 to 1) compares these cophenetic distances with the original assignment probabilities, providing an overall measure of clustering stability.

In our analyses, both stability metrics peaked at K = 3 (**Figure S5**), which was therefore selected for downstream analyses.

### Determining the most representative random state

We identified the most representative state from 100 random states (K = 3) using two key metrics: cosine similarity and archetype distinctiveness. Cosine similarity was used to assess how closely each state’s archetypes aligned with the mean archetype configuration, ensuring consistency. Simultaneously, we measured the distinctiveness of archetypes by measuring the euclidean distances between the 3 archetypes in each random state, to uphold the core principle of archetypal analysis—identifying extreme points. Our final scoring formula combined similarity to the mean with half-weighted distinctiveness, striking a balance between avoiding outlier states and maintaining representativeness. The selected state is visualized in the K-means clustering plot **(Figure S6)**, highlighting its optimal fit for subsequent analyses.

### Pathways’ compounds visualization using graphs

Network graphs were created to represent pathway interactions. The input data was sourced from the MetaCyc database^66^ and consisted of edges annotated with pathway-specific metabolites. Nodes corresponding to compounds shared across multiple pathways were connected using dashed gray edges to highlight duplication and identify clusters of shared compounds. Common compounds, such as H+, phosphate, ATP, ADP, H2O, NADP+, NADPH, NADH, NAD+, CO2, coenzyme A, AMP, dioxygen, hydrogen carbonate, and diphosphate, were excluded to reduce visual noise. A small subset of pathways did not have graph data available and therefore were not included in the visualization - this was the case for 4 pathways in the top 20 defining archetype 2: PRPP-PWY: superpathway of histidine, purine, and pyrimidine biosynthesis; TCA-GLYOX-BYPASS: superpathway of glyoxylate bypass and TCA; TCA: TCA cycle I (prokaryotic); METSYN-PWY: superpathway of L-homoserine and L-methionine biosynthesis; however they often had other highly similar pathways presented in the graph.

The finalized graphs were exported as HTML files, allowing interactive exploration and further visual refinement.

### Archetype stability analysis

To assess the stability of the archetypes, we filtered our dataset to identify healthy subjects with at least two unique samples, which yielded seven studies (n = 656 subjects, 1,557 samples). These subjects had 2–6 visits over a 2–730 day span (**Table S3**). We subtracted the archetype values between consecutive visits to calculate differences. For analyses based on archetype usage, samples were categorized as high usage (≥ 0.66) or medium usage (0.33–0.66).

### Differential analysis of disease samples using MaAsLin2

We curated metagenomic profiles from studies of patients with inflammatory bowel disease (IBD; 3 studies), type 2 diabetes (T2D; 3 studies), and colorectal cancer (CRC; 5 studies) (**Table S4A).** We then applied MaAsLin2^67^ to identify significant functional differential relative abundances in gut microbiomes between healthy individuals and those with inflammatory bowel disease (IBD), type 2 diabetes (T2D) and colorectal cancer (CRC) samples, respectively. All analyses were performed using the negative binomial model to address the compositional nature of the microbiome and account for overdispersion in the data and cumulative sum scaling normalization to adjust for library size, and Benjamini-Hochberg FDR to adjusted for multiple testing, as implemented in MaAslin2 R/Bioconductor (V1.18.0). In all analyses, subject ID and study were modeled as random effects to account for inter-individual variability and differences between datasets, respectively.

We conducted the following analyses:

1. Traditional analysis without archetype adjustment: Disease status (disease vs. healthy) was included as the fixed effect without any adjustment for archetypes.
2. Analyses with archetype adjustment: These models extended the “traditional” analysis by adding three fixed-effect to adjust for samples’ archetype usage (archetype1, archetype2, and archetype3)
3. Stratified analyses with archetype adjustment: Samples were stratified based on their dominant archetype (highest archetype value), and each stratified group was analyzed separately. For each group, the model included disease status as a fixed effect along with archetype usage values for archetype1, archetype2, and archetype3.

## References

1. Ley, R.E., Lozupone, C.A., Hamady, M., Knight, R., and Gordon, J.I. (2008). Worlds within worlds: evolution of the vertebrate gut microbiota. Nat. Rev. Microbiol. 6, 776–788. 10.1038/nrmicro1978.

2. Tropini, C., Earle, K.A., Huang, K.C., and Sonnenburg, J.L. (2017). The Gut Microbiome: Connecting Spatial Organization to Function. Cell Host Microbe 21, 433–442. 10.1016/j.chom.2017.03.010.

3. Conwill, A., Kuan, A.C., Damerla, R., Poret, A.J., Baker, J.S., Tripp, A.D., Alm, E.J., and Lieberman, T.D. (2022). Anatomy promotes neutral coexistence of strains in the human skin microbiome. Cell Host Microbe 30, 171–182.e7. 10.1016/j.chom.2021.12.007.

4. Coyte, K.Z., Schluter, J., and Foster, K.R. (2015). The ecology of the microbiome: Networks, competition, and stability. Science 350, 663–666. 10.1126/science.aad2602.

5. Heintz-Buschart, A., and Wilmes, P. (2018). Human Gut Microbiome: Function Matters. Trends Microbiol. 26, 563–574. 10.1016/j.tim.2017.11.002.

6. Hu, J., Amor, D.R., Barbier, M., Bunin, G., and Gore, J. (2022). Emergent phases of ecological diversity and dynamics mapped in microcosms. Science 378, 85–89. 10.1126/science.abm7841.

7. McKinlay, J.B. (2023). Are Bacteria Leaky? Mechanisms of Metabolite Externalization in Bacterial Cross-Feeding. Annu. Rev. Microbiol. 77, null. 10.1146/annurev-micro-032521-023815.

8. Seth, E.C., and Taga, M.E. (2014). Nutrient cross-feeding in the microbial world. Front. Microbiol. 5.

9. Hibbing, M.E., Fuqua, C., Parsek, M.R., and Peterson, S.B. (2010). Bacterial competition: surviving and thriving in the microbial jungle. Nat. Rev. Microbiol. 8, 15–25. 10.1038/nrmicro2259.

10. Rothschild, D., Leviatan, S., Hanemann, A., Cohen, Y., Weissbrod, O., and Segal, E. (2022). An atlas of robust microbiome associations with phenotypic traits based on large-scale cohorts from two continents. PLOS ONE 17, e0265756. 10.1371/journal.pone.0265756.

11. Costello, E.K., Lauber, C.L., Hamady, M., Fierer, N., Gordon, J.I., and Knight, R. (2009). Bacterial Community Variation in Human Body Habitats Across Space and Time. Science 326, 1694–1697. 10.1126/science.1177486.

12. Shenhav, L., Furman, O., Briscoe, L., Thompson, M., Silverman, J.D., Mizrahi, I., and Halperin, E. (2019). Modeling the temporal dynamics of the gut microbial community in adults and infants. PLOS Comput. Biol. 15, e1006960. 10.1371/journal.pcbi.1006960.

13. Arumugam, M., Raes, J., Pelletier, E., Le Paslier, D., Yamada, T., Mende, D.R., Fernandes, G.R., Tap, J., Bruls, T., Batto, J.-M., et al. (2011). Enterotypes of the human gut microbiome. Nature 473, 174–180. 10.1038/nature09944.

14. Gibson, T.E., Bashan, A., Cao, H.-T., Weiss, S.T., and Liu, Y.-Y. (2016). On the Origins and Control of Community Types in the Human Microbiome. PLoS Comput. Biol. 12, e1004688. 10.1371/journal.pcbi.1004688.

15. Human Microbiome Project Consortium (2012). Structure, function and diversity of the healthy human microbiome. Nature 486, 207–214. 10.1038/nature11234.

16. Yatsunenko, T., Rey, F.E., Manary, M.J., Trehan, I., Dominguez-Bello, M.G., Contreras, M., Magris, M., Hidalgo, G., Baldassano, R.N., Anokhin, A.P., et al. (2012). Human gut microbiome viewed across age and geography. Nature 486, 222–227. 10.1038/nature11053.

17. Costea, P.I., Hildebrand, F., Arumugam, M., Bäckhed, F., Blaser, M.J., Bushman, F.D., de Vos, W.M., Ehrlich, S.D., Fraser, C.M., Hattori, M., et al. (2018). Enterotypes in the landscape of gut microbial community composition. Nat. Microbiol. 3, 8–16. 10.1038/s41564-017-0072-8.

18. Wu, G.D., Chen, J., Hoffmann, C., Bittinger, K., Chen, Y.-Y., Keilbaugh, S.A., Bewtra, M., Knights, D., Walters, W.A., Knight, R., et al. (2011). Linking long-term dietary patterns with gut microbial enterotypes. Science 334, 105–108. 10.1126/science.1208344.

19. Ding, T., and Schloss, P.D. (2014). Dynamics and associations of microbial community types across the human body. Nature 509, 357–360. 10.1038/nature13178.

20. Hildebrand, F., Nguyen, T.L.A., Brinkman, B., Yunta, R.G., Cauwe, B., Vandenabeele, P., Liston, A., and Raes, J. (2013). Inflammation-associated enterotypes, host genotype, cage and inter-individual effects drive gut microbiota variation in common laboratory mice. Genome Biol. 14, R4. 10.1186/gb-2013-14-1-r4.

21. Moeller, A.H., Degnan, P.H., Pusey, A.E., Wilson, M.L., Hahn, B.H., and Ochman, H. (2012). Chimpanzees and humans harbour compositionally similar gut enterotypes. Nat. Commun. 3, 1179. 10.1038/ncomms2159.

22. Zhou, Y., Mihindukulasuriya, K.A., Gao, H., La Rosa, P.S., Wylie, K.M., Martin, J.C., Kota, K., Shannon, W.D., Mitreva, M., Sodergren, E., et al. (2014). Exploration of bacterial community classes in major human habitats. Genome Biol. 15, R66. 10.1186/gb-2014-15-5-r66.

23. Frioux, C., Ansorge, R., Özkurt, E., Nedjad, C.G., Fritscher, J., Quince, C., Waszak, S.M., and Hildebrand, F. (2023). Enterosignatures define common bacterial guilds in the human gut microbiome. Cell Host Microbe 31, 1111–1125.e6. 10.1016/j.chom.2023.05.024.

24. Keller, M.I., Nishijima, S., Podlesny, D., Kim, C.Y., Robbani, S.M., Schudoma, C., Fullam, A., Richter, J., Letunic, I., Akanni, W., et al. (2024). Refined Enterotyping Reveals Dysbiosis in Global Fecal Metagenomes. Preprint at bioRxiv, https://doi.org/10.1101/2024.08.13.607711 10.1101/2024.08.13.607711.

25. Knights, D., Ward, T.L., McKinlay, C.E., Miller, H., Gonzalez, A., McDonald, D., and Knight, R. (2014). Rethinking “Enterotypes.” Cell Host Microbe 16, 433–437. 10.1016/j.chom.2014.09.013.

26. Levy, R., Magis, A.T., Earls, J.C., Manor, O., Wilmanski, T., Lovejoy, J., Gibbons, S.M., Omenn, G.S., Hood, L., and Price, N.D. (2020). Longitudinal analysis reveals transition barriers between dominant ecological states in the gut microbiome. Proc. Natl. Acad. Sci. 117, 13839–13845. 10.1073/pnas.1922498117.

27. Reichardt, N., Vollmer, M., Holtrop, G., Farquharson, F.M., Wefers, D., Bunzel, M., Duncan, S.H., Drew, J.E., Williams, L.M., Milligan, G., et al. (2018). Specific substrate-driven changes in human faecal microbiota composition contrast with functional redundancy in short-chain fatty acid production. ISME J. 12, 610–622. 10.1038/ismej.2017.196.

28. Watson, A.R., Füssel, J., Veseli, I., DeLongchamp, J.Z., Silva, M., Trigodet, F., Lolans, K., Shaiber, A., Fogarty, E., Runde, J.M., et al. (2023). Metabolic independence drives gut microbial colonization and resilience in health and disease. Genome Biol. 24, 78. 10.1186/s13059-023-02924-x.

29. Cutler, A., and Breiman, L. (1994). Archetypal Analysis. Technometrics 36, 338–347. 10.1080/00401706.1994.10485840.

30. Keller, S.M., Samarin, M., Arend Torres, F., Wieser, M., and Roth, V. (2021). Learning Extremal Representations with Deep Archetypal Analysis. Int. J. Comput. Vis. 129, 805–820. 10.1007/s11263-020-01390-3.

31. van Dijk, D., Burkhardt, D., Amodio, M., Tong, A., Wolf, G., and Krishnaswamy, S. (2019). Finding Archetypal Spaces Using Neural Networks. ArXiv190109078 Cs Stat.

32. Wang, Y., and Zhao, H. (2022). Non-linear archetypal analysis of single-cell RNA-seq data by deep autoencoders. PLOS Comput. Biol. 18, e1010025. 10.1371/journal.pcbi.1010025.

33. Pasolli, E., Schiffer, L., Manghi, P., Renson, A., Obenchain, V., Truong, D.T., Beghini, F., Malik, F., Ramos, M., Dowd, J.B., et al. (2017). Accessible, curated metagenomic data through ExperimentHub. Nat. Methods 14, 1023–1024. 10.1038/nmeth.4468.

34. Dai, D., Zhu, J., Sun, C., Li, M., Liu, J., Wu, S., Ning, K., He, L., Zhao, X.-M., and Chen, W.-H. (2021). GMrepo v2: a curated human gut microbiome database with special focus on disease markers and cross-dataset comparison. Nucleic Acids Res. 50, D777–D784. 10.1093/nar/gkab1019.

35. Tigchelaar, E.F., Zhernakova, A., Dekens, J.A.M., Hermes, G., Baranska, A., Mujagic, Z., Swertz, M.A., Muñoz, A.M., Deelen, P., Cénit, M.C., et al. (2015). Cohort profile: LifeLines DEEP, a prospective, general population cohort study in the northern Netherlands: study design and baseline characteristics. BMJ Open 5, e006772. 10.1136/bmjopen-2014-006772.

36. Scepanovic, P., Hodel, F., Mondot, S., Partula, V., Byrd, A., Hammer, C., Alanio, C., Bergstedt, J., Patin, E., Touvier, M., et al. (2019). A comprehensive assessment of demographic, environmental, and host genetic associations with gut microbiome diversity in healthy individuals. Microbiome 7, 130. 10.1186/s40168-019-0747-x.

37. Zhang, Y., Parmigiani, G., and Johnson, W.E. (2020). ComBat-seq: batch effect adjustment for RNA-seq count data. NAR Genomics Bioinforma. 2, lqaa078. 10.1093/nargab/lqaa078.

38. Zhu, W., Winter, M.G., Byndloss, M.X., Spiga, L., Duerkop, B.A., Hughes, E.R., Büttner, L., de Lima Romão, E., Behrendt, C.L., Lopez, C.A., et al. (2018). Precision editing of the gut microbiota ameliorates colitis. Nature 553, 208–211. 10.1038/nature25172.

39. De Filippis, F., Pasolli, E., Tett, A., Tarallo, S., Naccarati, A., De Angelis, M., Neviani, E., Cocolin, L., Gobbetti, M., Segata, N., et al. (2019). Distinct Genetic and Functional Traits of Human Intestinal *Prevotella copri* Strains Are Associated with Different Habitual Diets. Cell Host Microbe 25, 444–453.e3. 10.1016/j.chom.2019.01.004.

40. Connors, J., Dawe, N., and Van Limbergen, J. (2018). The Role of Succinate in the Regulation of Intestinal Inflammation. Nutrients 11, 25. 10.3390/nu11010025.

41. Fernández-Veledo, S., and Vendrell, J. (2019). Gut microbiota-derived succinate: Friend or foe in human metabolic diseases? Rev. Endocr. Metab. Disord. 20, 439–447. 10.1007/s11154-019-09513-z.

42. Garcia-Mazcorro, J.F., Amieva-Balmori, M., Triana-Romero, A., Wilson, B., Smith, L., Reyes-Huerta, J., Rossi, M., Whelan, K., and Remes-Troche, J.M. (2023). Fecal Microbial Composition and Predicted Functional Profile in Irritable Bowel Syndrome Differ between Subtypes and Geographical Locations. Microorganisms 11, 2493. 10.3390/microorganisms11102493.

43. Gargari, G., Mantegazza, G., Taverniti, V., Gardana, C., Valenza, A., Rossignoli, F., Barbaro, M.R., Marasco, G., Cremon, C., Barbara, G., et al. (2023). Fecal short-chain fatty acids in non-constipated irritable bowel syndrome: a potential clinically relevant stratification factor based on catabotyping analysis. Gut Microbes 15, 2274128. 10.1080/19490976.2023.2274128.

44. Su, Q., Tun, H.M., Liu, Q., Yeoh, Y.K., Mak, J.W.Y., Chan, F.K., and Ng, S.C. (2023). Gut microbiome signatures reflect different subtypes of irritable bowel syndrome. Gut Microbes 15, 2157697. 10.1080/19490976.2022.2157697.

45. Vervier, K., Moss, S., Kumar, N., Adoum, A., Barne, M., Browne, H., Kaser, A., Kiely, C.J., Neville, B.A., Powell, N., et al. (2022). Two microbiota subtypes identified in irritable bowel syndrome with distinct responses to the low FODMAP diet. Gut 71, 1821–1830. 10.1136/gutjnl-2021-325177.

46. Chen, C., Yan, W., Tao, M., and Fu, Y. (2023). NAD+ Metabolism and Immune Regulation: New Approaches to Inflammatory Bowel Disease Therapies. Antioxidants 12, 1230. 10.3390/antiox12061230.

47. Gerner, R.R., Klepsch, V., Macheiner, S., Arnhard, K., Adolph, T.E., Grander, C., Wieser, V., Pfister, A., Moser, P., Hermann-Kleiter, N., et al. (2018). NAD metabolism fuels human and mouse intestinal inflammation. Gut 67, 1813–1823. 10.1136/gutjnl-2017-314241.

48. Li, P., Yin, Y.-L., Li, D., Kim, S.W., and Wu, G. (2007). Amino acids and immune function. Br. J. Nutr. 98, 237–252. 10.1017/S000711450769936X.

49. Liu, Y., Wang, X., Hou, Y., Yin, Y., Qiu, Y., Wu, G., and Hu, C.-A.A. (2017). Roles of amino acids in preventing and treating intestinal diseases: recent studies with pig models. Amino Acids 49, 1277–1291. 10.1007/s00726-017-2450-1.

50. Ruth, M.R., and Field, C.J. (2013). The immune modifying effects of amino acids on gut-associated lymphoid tissue. J. Anim. Sci. Biotechnol. 4, 27. 10.1186/2049-1891-4-27.

51. Kittana, H., Gomes-Neto, J.C., Heck, K., Juritsch, A.F., Sughroue, J., Xian, Y., Mantz, S., Segura Muñoz, R.R., Cody, L.A., Schmaltz, R.J., et al. (2023). Evidence for a Causal Role for Escherichia coli Strains Identified as Adherent-Invasive (AIEC) in Intestinal Inflammation. mSphere 8, e00478–22. 10.1128/msphere.00478-22.

52. Palmela, C., Chevarin, C., Xu, Z., Torres, J., Sevrin, G., Hirten, R., Barnich, N., Ng, S.C., and Colombel, J.-F. (2018). Adherent-invasive Escherichia coli in inflammatory bowel disease. Gut 67, 574–587. 10.1136/gutjnl-2017-314903.

53. Yin, R., Wang, T., Dai, H., Han, J., Sun, J., Liu, N., Dong, W., Zhong, J., and Liu, H. (2023). Immunogenic molecules associated with gut bacterial cell walls: chemical structures, immune-modulating functions, and mechanisms. Protein Cell 14, 776–785. 10.1093/procel/pwad016.

54. Chen, S., Zhou, Y., Chen, Y., and Gu, J. (2018). fastp: an ultra-fast all-in-one FASTQ preprocessor. Bioinformatics 34, i884–i890. 10.1093/bioinformatics/bty560.

55. Simpson, S., Bettauer, V., Ramachandran, A., Kraemer, S., Mahon, S., Medina, M., Vallès, Y., Dumeaux, V., Vallès, H., Walsh, D., et al. (2023). A metagenomic-based study of two sites from the Barbadian reef system. Coral Reefs Online 42, 359–366. 10.1007/s00338-022-02330-y.

56. Wright, R.J., Comeau, A.M., and Langille, M.G.I. (2023). From defaults to databases: parameter and database choice dramatically impact the performance of metagenomic taxonomic classification tools. Microb. Genomics 9, 000949. 10.1099/mgen.0.000949.

57. Ye, S.H., Siddle, K.J., Park, D.J., and Sabeti, P.C. (2019). Benchmarking Metagenomics Tools for Taxonomic Classification. Cell 178, 779–794. 10.1016/j.cell.2019.07.010.

58. Lu, J., Breitwieser, F.P., Thielen, P., and Salzberg, S.L. (2017). Bracken: estimating species abundance in metagenomics data. PeerJ Comput. Sci. 3, e104. 10.7717/peerj-cs.104.

59. Wood, D.E., Lu, J., and Langmead, B. (2019). Improved metagenomic analysis with Kraken 2. Genome Biol. 20, 257. 10.1186/s13059-019-1891-0.

60. Hiseni, P., Rudi, K., Wilson, R.C., Hegge, F.T., and Snipen, L. (2021). HumGut: a comprehensive human gut prokaryotic genomes collection filtered by metagenome data. Microbiome 9, 165. 10.1186/s40168-021-01114-w.

61. Parks, D.H., Chuvochina, M., Rinke, C., Mussig, A.J., Chaumeil, P.-A., and Hugenholtz, P. (2022). GTDB: an ongoing census of bacterial and archaeal diversity through a phylogenetically consistent, rank normalized and complete genome-based taxonomy. Nucleic Acids Res. 50, D785–D794. 10.1093/nar/gkab776.

62. Beghini, F., McIver, L.J., Blanco-Míguez, A., Dubois, L., Asnicar, F., Maharjan, S., Mailyan, A., Manghi, P., Scholz, M., Thomas, A.M., et al. (2021). Integrating taxonomic, functional, and strain-level profiling of diverse microbial communities with bioBakery 3. eLife 10, e65088. 10.7554/eLife.65088.

63. de la Cuesta-Zuluaga, J., Ley, R.E., and Youngblut, N.D. (2020). Struo: a pipeline for building custom databases for common metagenome profilers. Bioinformatics 36, 2314–2315. 10.1093/bioinformatics/btz899.

64. Song, D., Wang, Q., Yan, G., Liu, T., Sun, T., and Li, J.J. (2023). scDesign3 generates realistic in silico data for multimodal single-cell and spatial omics. Nat. Biotechnol. 10.1038/s41587-023-01772-1.

65. Wang, Y., and Zhao, H. (2022). Non-linear archetypal analysis of single-cell RNA-seq data by deep autoencoders. PLOS Comput. Biol. 18, e1010025. 10.1371/journal.pcbi.1010025.

66. Caspi, R., Altman, T., Billington, R., Dreher, K., Foerster, H., Fulcher, C.A., Holland, T.A., Keseler, I.M., Kothari, A., Kubo, A., et al. (2014). The MetaCyc database of metabolic pathways and enzymes and the BioCyc collection of Pathway/Genome Databases. Nucleic Acids Res. 42, D459–D471. 10.1093/nar/gkt1103.

67. Mallick, H., Rahnavard, A., and McIver, L. (2022). Maaslin2: “Multivariable Association Discovery in Population-scale Meta-omics Studies.” Version 1.10.0 (Bioconductor version: Release (3.15)). https://doi.org/10.18129/B9.bioc.Maaslin2 10.18129/B9.bioc.Maaslin2.

