## Supplemental Information for "Deep learning reveals functional archetypes in the adult human gut microbiome that underlie interindividual variability and confound disease signals"

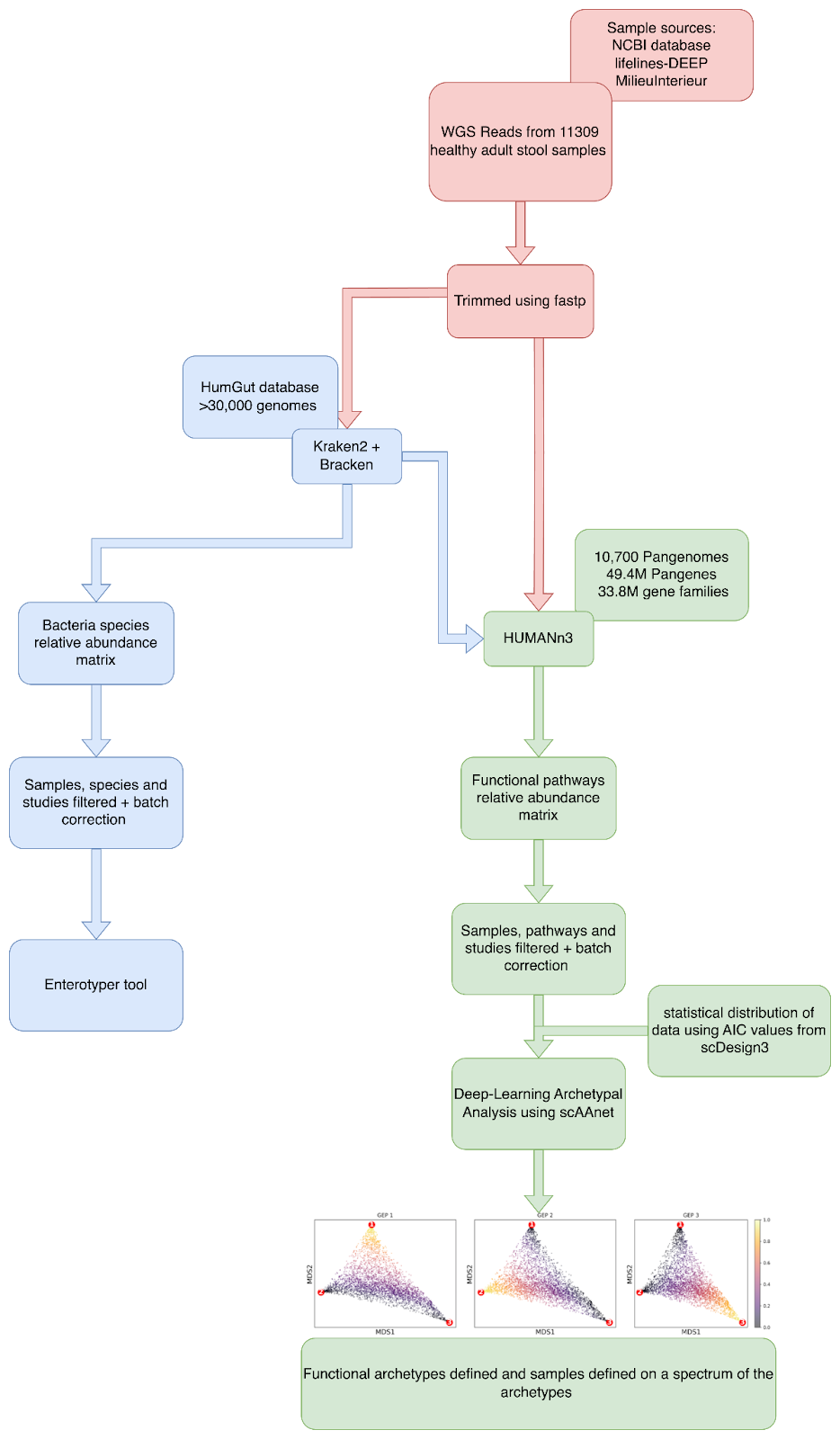

**Figure S1.** Illustration of the data analysis pipeline and summary of methods used.

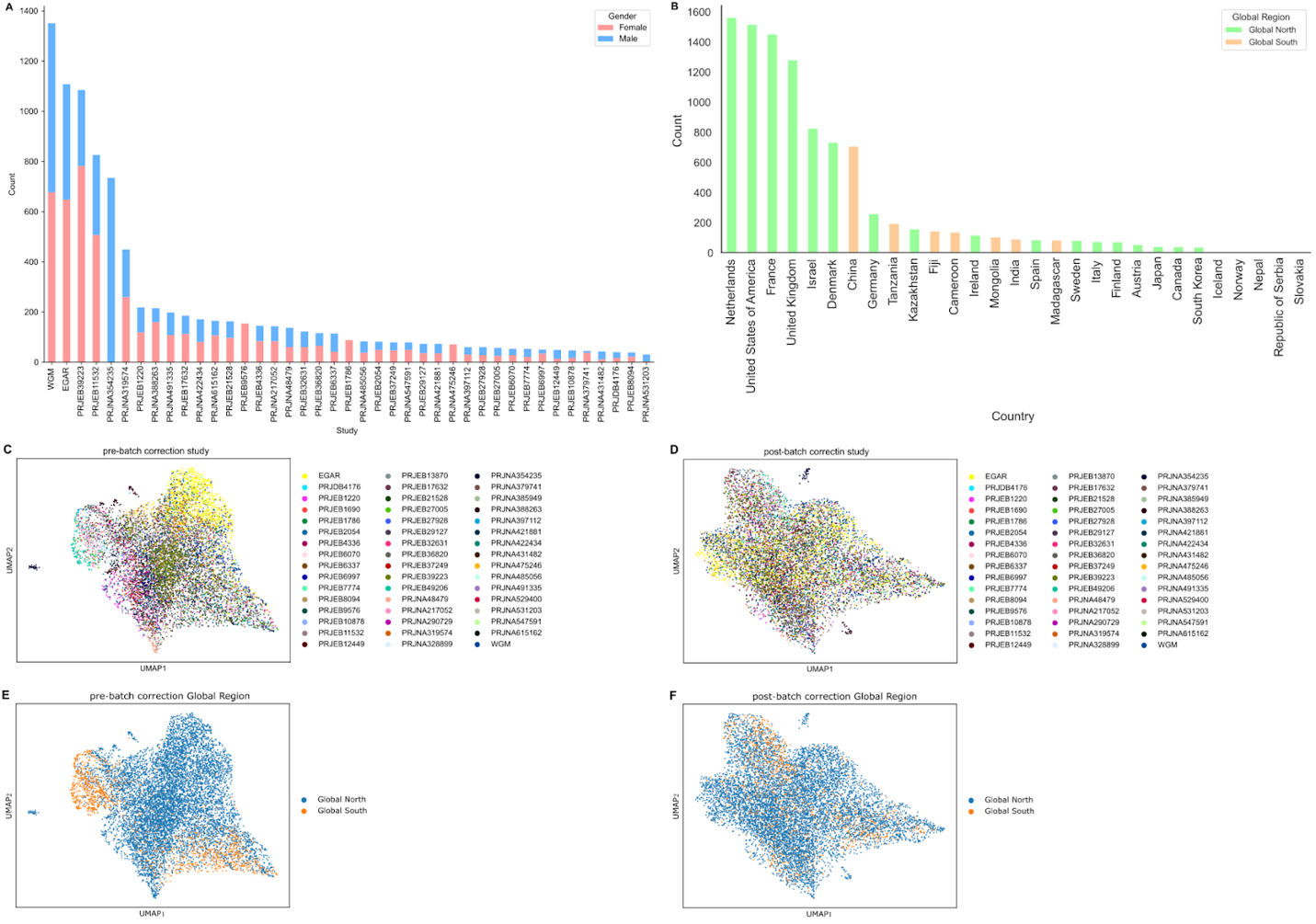

**Figure S2. (A, B)** Bar plots of samples that passed the following filtering criteria: total read counts (library sizes) between 100,000 and 2,000,000, and removal of studies with fewer than 30 samples. In **(A)**, bars are grouped by study and colored by gender; in **(B)**, bars are grouped by country and colored by global region. **(C–F)** UMAP projections of samples before and after the above filtering and batch correction using ComBat-seq. In **(C, D)**, samples are colored by study source, while in **(E, F)**, samples are colored by global region.

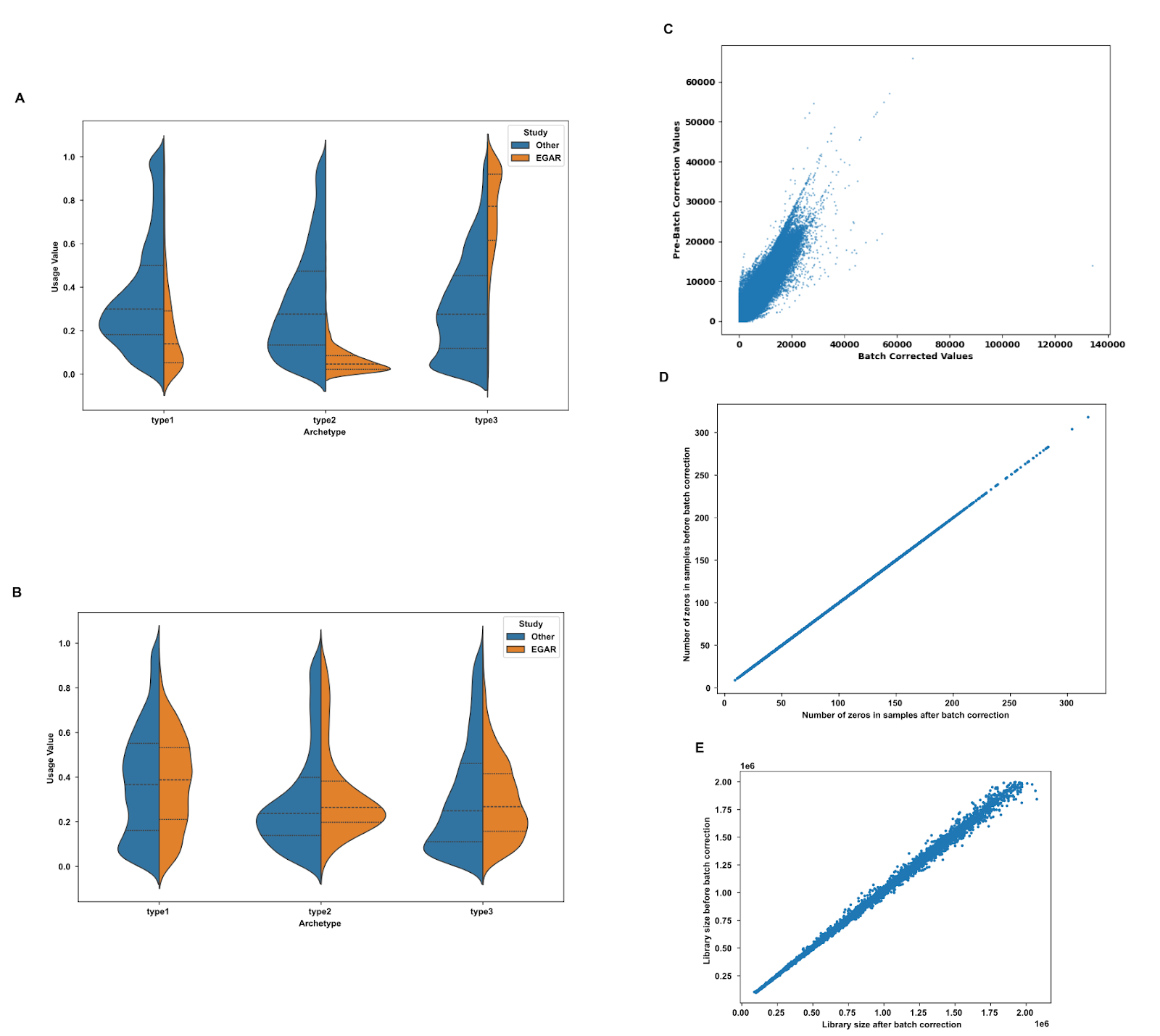

**Figure S3.** Violin plots comparing the assigned archetype values for the study that showed the largest batch effect to samples from other studies **(A)** before batch correction **(B)** after batch correction. **(C-E)** Scatter plots comparing values before and after batch correction **(C)** of each pathway count in each sample **(D)** The sparsity (zero counts) of samples **(E)** The library size of samples.

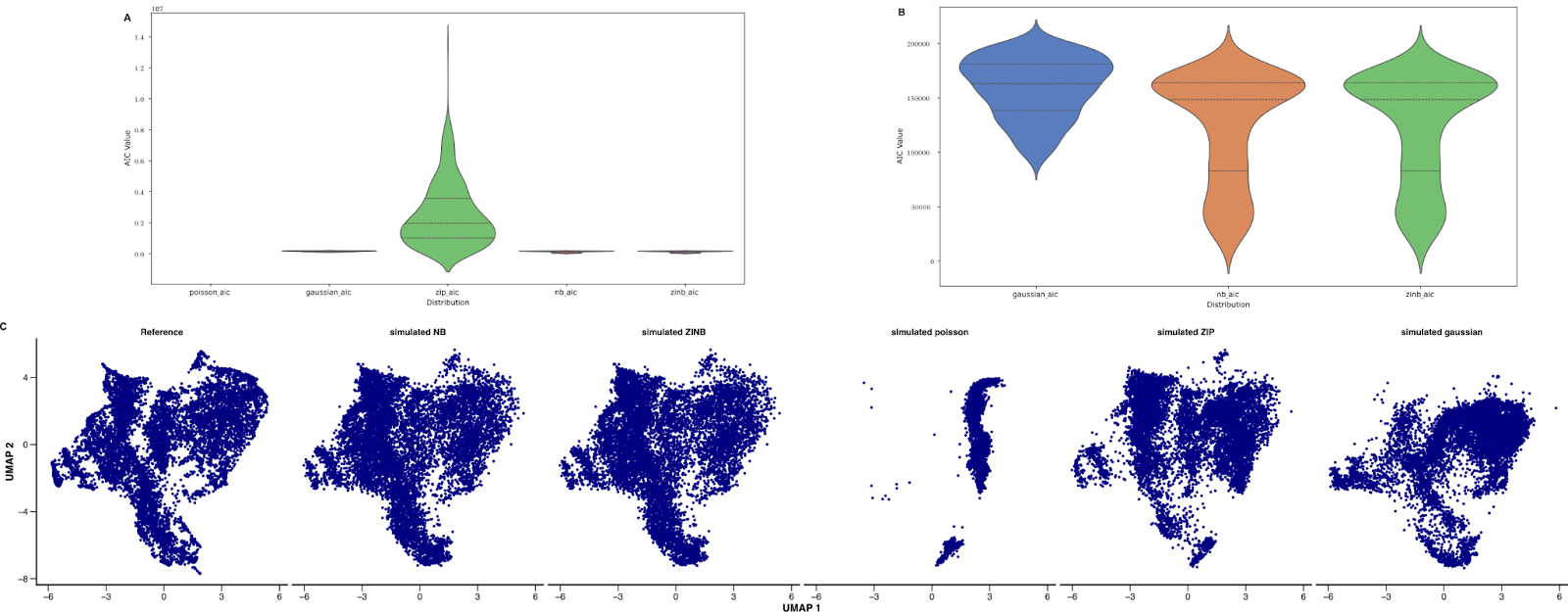

**Figure S4. (A, B)** Violin plots showing the distribution of Akaike information criterion (AIC) values of **(A)** poisson, zero inflated poisson (ZIP), gaussian, negative binomial(NB) and zero inflated negative binomial (ZINB). Poisson values were infinite and hence do not appear on the violin plot. **(B)** distribution of gaussian, negative binomial(NB) and zero inflated negative binomial (ZINB) only to avoid the high AIC values of ZIP from affecting visualization. **(C)** UMAP comparing sample simulation performance when assuming functional pathway data follows negative binomial (NB), zero inflated negative binomial (ZINB), poisson, zero inflated poisson (ZIP) and gaussian distributions.

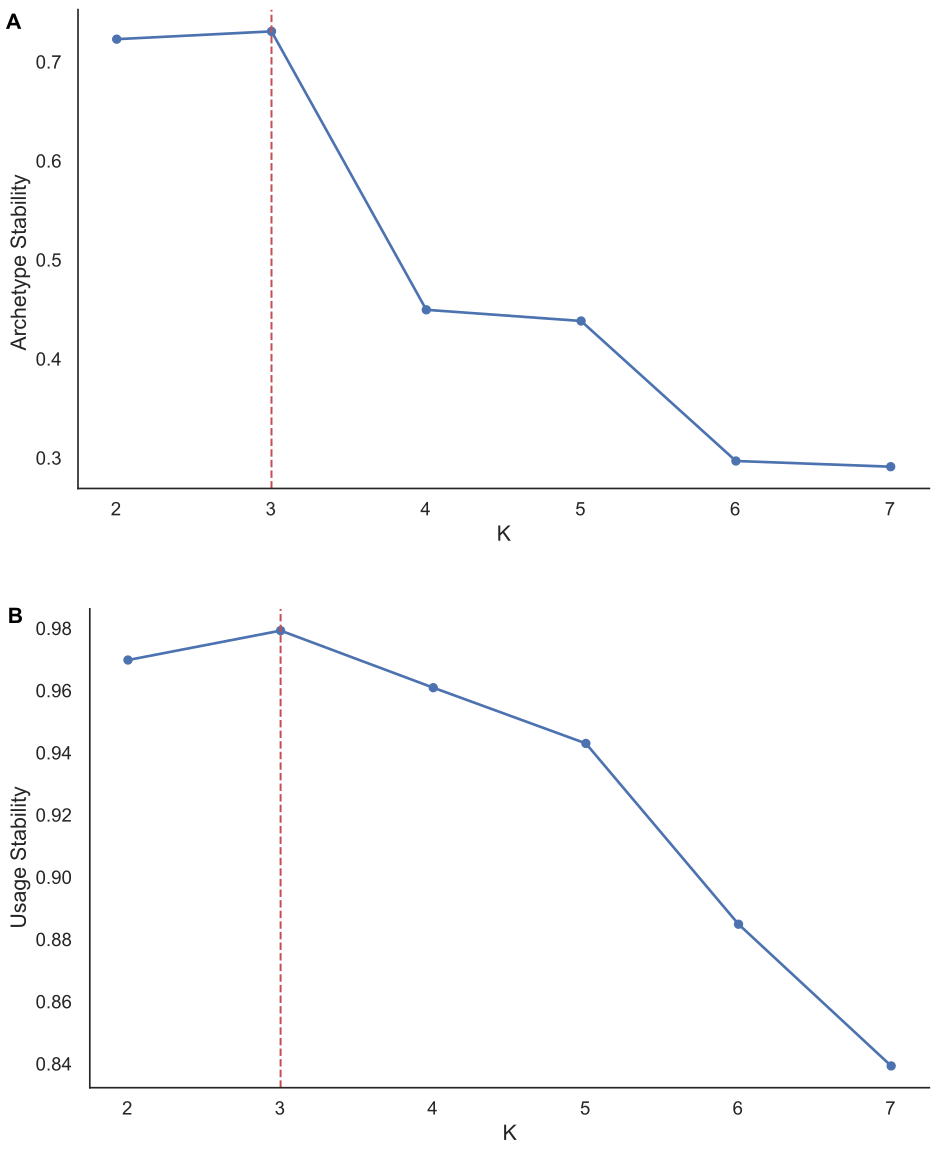

**Figure S5.** Model stability results after running scAAnet across K values of 2 to 7 over 8 random states. **(A)** Archetypes’ stability results and **(B)** Samples’ archetype usage stability results. Red dashed line indicates the final value of K = 3 was selected for further analysis. **(C)** Table of archetypes’ stability values and usage stability values after running scAAnet across K values of 2 to 7 over 8 random states.

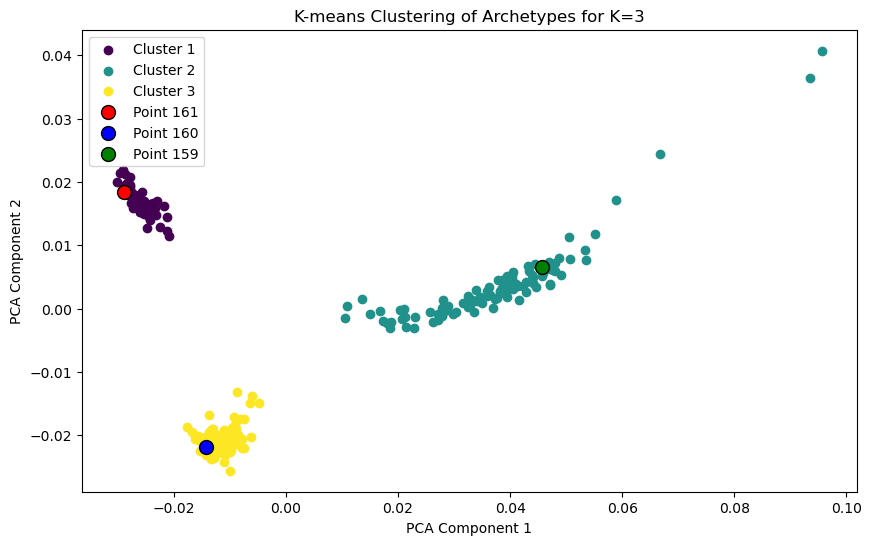

**Figure S6.** PCA plot with K means clustering of the archetypes’ from 100 random states. Each state had 3 archetypes, giving a total of 300 points, each cluster (each representing an archetype) having 100 points. Points 159, 160 and 161 (red, blue and green labeled points) are the 3 archetypes from state number 53, which was selected as the most representative state based on the mean of all the states and how different its own 3 archetypes are.

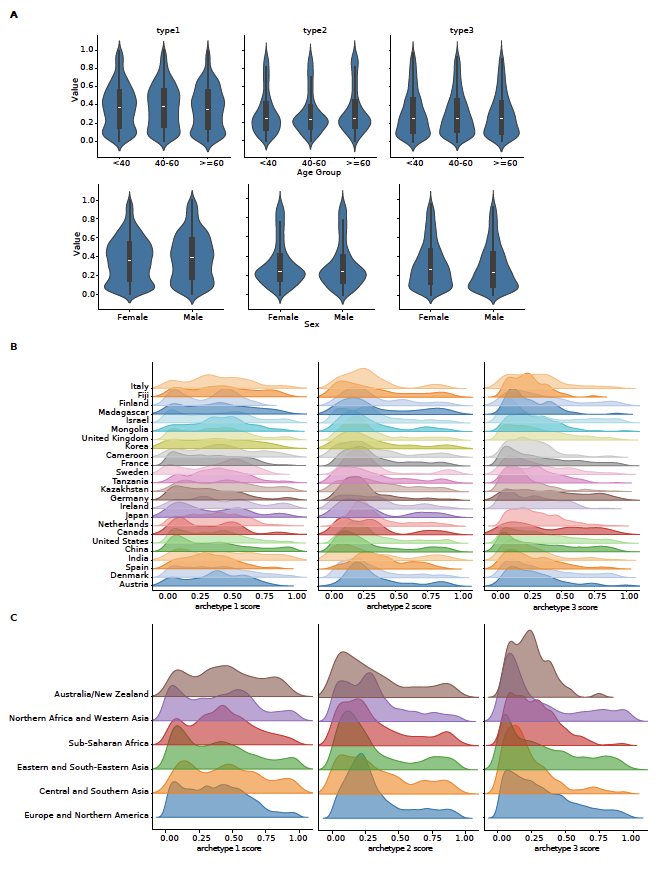

**Figure S7.** Distribution for archetype 1 (left), archetype 2 (middle) and archetype 3 (right) scores across **(A)** age groups (top) and sex (bottom), **(B)** country, or **(C)** world region of origin.

| **Repository** | **BioPorject Accession** |
| --- | --- |
| GMrepo V2 | PRJNA397219^1^  PRJEB7774^2^  PRJDB4176^3^  PRJEB6997^4^  PRJNA422434^5^  PRJEB15371^6^  PRJNA388263^7^  PRJEB2054^8^  PRJEB1786^9^  PRJNA298489^10^  PRJNA615162^11^  PRJEB8094^12^  PRJNA447983^13^  PRJNA428202^14^  PRJNA389280^15^  PRJNA373879^16^  PRJEB10878^17^  PRJEB6070^18^ |
| CuratedMetagenomicData | PRJNA504891^19^  PRJEB1220^20^  PRJNA278393^21^  PRJNA397112^22^  PRJNA379120^23^  PRJNA431482^24^  PRJEB12449^25^  PRJEB17784^26^  PRJEB6337^27^  PRJNA491335^28^  PRJEB38742^29^  PRJEB36820^30^  PRJEB27928^31^  PRJEB17632^32^  PRJNA529400^33^  PRJNA354235^34^  PRJNA547591^35^  PRJEB1690^36^  PRJEB4336^37^  PRJNA529124^33^  PRJEB47909^38^  PRJNA389927^39^  PRJEB32631^40^  PRJEB9150^41^  PRJNA290729^42^  PRJNA319574^43^  PRJEB29127^44^  PRJNA328899^45^  PRJNA289586^46^  PRJEB9576^47^  PRJNA392180^48^  PRJNA379741^49^  PRJNA352475^50^  PRJEB6456^51^  PRJEB11532^52^  PRJEB7369^53^  PRJNA485056^19^  PRJNA299502^54^  PRJEB13870^55^  PRJEB21528^56^  PRJEB27005^57^  PRJNA339914^58^  PRJNA48479^59^  PRJNA475246^60^  PRJNA217052^61^  PRJEB18780^62^  PRJEB37249^63^  PRJNA53120^64^  PRJEB12357^65^  PRJNA340216^66^  PRJNA385949^67^  PRJNA407815^68^  PRJNA297252^69^  PRJNA421881^70^ |

**Table S1.** List of studies included in this study curated from two dataset repositories: GMrepo and CuratedMetagenomicData R package. Full references are listed at the end of this document.

| **Color** | **Group** |
| --- | --- |
|  | Sugars metabolism |
|  | BCAA biosynthesis |
|  | Peptidoglycan and Cell Wall Biosynthesis |
|  | Fatty acids metabolism |
|  | Amino acids and its derivatives synthesis and degradation |
|  | Coenzyme Biosynthesis (Used in amino acid metabolism) |
|  | Methionine synthesis |
|  | Nucleotide synthesis |
|  | Other specialized pathways |

| **Pathway** | **Group** | **Type1** | **Type2** | **Type3** |
| --- | --- | --- | --- | --- |
| DTDPRHAMSYN-PWY: dTDP-&beta;-L-rhamnose biosynthesis |  | **1.00** | 0.00 | 0.39 |
| PWY-7237: myo-, chiro- and scyllo-inositol degradation |  | **0.73** | 0.06 | 0.10 |
| GLUCOSE1PMETAB-PWY: glucose and glucose-1-phosphate degradation |  | **0.52** | 0.10 | 0.00 |
| PWY-7221: guanosine ribonucleotides de novo biosynthesis |  | **0.48** | 0.03 | 0.18 |
| PWY-1042: glycolysis IV |  | **0.48** | 0.03 | 0.06 |
| VALSYN-PWY: L-valine biosynthesis |  | **0.45** | 0.03 | 0.15 |
| ILEUSYN-PWY: L-isoleucine biosynthesis I (from threonine) |  | **0.43** | 0.03 | 0.14 |
| BRANCHED-CHAIN-AA-SYN-PWY: superpathway of branched chain amino acid biosynthesis |  | **0.42** | 0.03 | 0.17 |
| PWY-5103: L-isoleucine biosynthesis III |  | **0.42** | 0.04 | 0.14 |
| PEPTIDOGLYCANSYN-PWY: peptidoglycan biosynthesis I (meso-diaminopimelate containing) |  | **0.37** | 0.04 | 0.14 |
| PWY-6387: UDP-N-acetylmuramoyl-pentapeptide biosynthesis I (meso-diaminopimelate containing) |  | **0.37** | 0.04 | 0.14 |
| PWY-6386: UDP-N-acetylmuramoyl-pentapeptide biosynthesis II (lysine-containing) |  | **0.35** | 0.05 | 0.14 |
| PWY-6151: S-adenosyl-L-methionine salvage I |  | **0.35** | 0.05 | 0.17 |
| PWY-7560: methylerythritol phosphate pathway II |  | **0.35** | 0.05 | 0.23 |
| PWY-6700: queuosine biosynthesis I (de novo) |  | **0.34** | 0.04 | 0.25 |
| PWY-5686: UMP biosynthesis I |  | **0.34** | 0.04 | 0.26 |
| PWY-7791: UMP biosynthesis III |  | **0.34** | 0.04 | 0.26 |
| PWY-7790: UMP biosynthesis II |  | **0.34** | 0.04 | 0.26 |
| PWY-6385: peptidoglycan biosynthesis III (mycobacteria) |  | **0.32** | 0.05 | 0.17 |
| PWY-7953: UDP-N-acetylmuramoyl-pentapeptide biosynthesis III (meso-diaminopimelate containing) |  | **0.32** | 0.05 | 0.15 |
| PRPP-PWY: superpathway of histidine, purine, and pyrimidine biosynthesis |  | 0.06 | **1.00** | 0.14 |
| DENOVOPURINE2-PWY: superpathway of purine nucleotides de novo biosynthesis II |  | 0.06 | **0.79** | 0.14 |
| TCA-GLYOX-BYPASS: superpathway of glyoxylate bypass and TCA |  | 0.06 | **0.76** | 0.14 |
| PWY-561: superpathway of glyoxylate cycle and fatty acid degradation |  | 0.06 | **0.54** | 0.14 |
| PWY-7187: pyrimidine deoxyribonucleotides de novo biosynthesis II |  | 0.06 | **0.54** | 0.14 |
| FAO-PWY: fatty acid &beta;-oxidation I (generic) |  | 0.06 | **0.53** | 0.14 |
| PWY0-1277: 3-phenylpropanoate and 3-(3-hydroxyphenyl)propanoate degradation |  | 0.06 | **0.51** | 0.14 |
| GLYCOLYSIS-TCA-GLYOX-BYPASS: superpathway of glycolysis, pyruvate dehydrogenase, TCA, and glyoxylate bypass |  | 0.06 | **0.43** | 0.14 |
| GLYOXYLATE-BYPASS: glyoxylate cycle |  | 0.06 | **0.42** | 0.13 |
| TCA: TCA cycle I (prokaryotic) |  | 0.06 | **0.41** | 0.13 |
| PWY-5675: nitrate reduction V (assimilatory) |  | 0.06 | **0.40** | 0.13 |
| PWY-6803: phosphatidylcholine acyl editing |  | 0.06 | **0.39** | 0.13 |
| AST-PWY: L-arginine degradation II (AST pathway) |  | 0.06 | **0.39** | 0.14 |
| PWY-5347: superpathway of L-methionine biosynthesis (transsulfuration) |  | 0.06 | **0.36** | 0.13 |
| MET-SAM-PWY: superpathway of S-adenosyl-L-methionine biosynthesis |  | 0.06 | **0.36** | 0.13 |
| METSYN-PWY: superpathway of L-homoserine and L-methionine biosynthesis |  | 0.06 | **0.35** | 0.13 |
| PWY-5136: fatty acid &beta;-oxidation II (plant peroxisome) |  | 0.06 | **0.34** | 0.12 |
| THREOCAT-PWY: superpathway of L-threonine metabolism |  | 0.06 | **0.34** | 0.14 |
| HCAMHPDEG-PWY: 3-phenylpropanoate and 3-(3-hydroxyphenyl)propanoate degradation to 2-hydroxypentadienoate |  | 0.06 | **0.33** | 0.14 |
| PWY-6690: cinnamate and 3-hydroxycinnamate degradation to 2-hydroxypentadienoate |  | 0.06 | **0.33** | 0.14 |
| PYRIDOXSYN-PWY: pyridoxal 5'-phosphate biosynthesis I |  | 0.05 | 0.11 | **1.00** |
| PWY-5030: L-histidine degradation III |  | 0.05 | 0.09 | **0.97** |
| ARGININE-SYN4-PWY: L-ornithine biosynthesis II |  | 0.03 | 0.09 | **0.93** |
| PWY-6902: chitin degradation II (Vibrio) |  | 0.05 | 0.11 | **0.90** |
| PWY0-845: superpathway of pyridoxal 5'-phosphate biosynthesis and salvage |  | 0.05 | 0.12 | **0.86** |
| HISDEG-PWY: L-histidine degradation I |  | 0.04 | 0.10 | **0.83** |
| PWY-7392: taxadiene biosynthesis (engineered) |  | 0.05 | 0.10 | **0.82** |
| PWY66-429: fatty acid biosynthesis initiation (mitochondria) |  | 0.00 | 0.07 | **0.78** |
| PWY-4984: urea cycle |  | 0.05 | 0.10 | **0.74** |
| PWY-6126: superpathway of adenosine nucleotides de novo biosynthesis II |  | 0.01 | 0.09 | **0.67** |
| PWY-7229: superpathway of adenosine nucleotides de novo biosynthesis I |  | 0.00 | 0.09 | **0.67** |
| CITRULBIO-PWY: L-citrulline biosynthesis |  | 0.04 | 0.11 | **0.64** |
| PWY-5130: 2-oxobutanoate degradation I |  | 0.05 | 0.10 | **0.64** |
| PWY-7228: superpathway of guanosine nucleotides de novo biosynthesis I |  | 0.01 | 0.09 | **0.64** |
| PWY-6125: superpathway of guanosine nucleotides de novo biosynthesis II |  | 0.01 | 0.09 | **0.62** |
| PWY-7220: adenosine deoxyribonucleotides de novo biosynthesis II |  | 0.02 | 0.10 | **0.62** |
| PWY-7222: guanosine deoxyribonucleotides de novo biosynthesis II |  | 0.02 | 0.10 | **0.62** |
| PWY-5121: superpathway of geranylgeranyl diphosphate biosynthesis II (via MEP) |  | 0.03 | 0.09 | **0.61** |
| PWY-5973: cis-vaccenate biosynthesis |  | 0.04 | 0.07 | **0.59** |
| PWY-7282: 4-amino-2-methyl-5-diphosphomethylpyrimidine biosynthesis II |  | 0.03 | 0.11 | **0.59** |

**Table S2.** Top 20 significant pathways and their relative importance values (scale of 0 to 1) for archetypes 1, 2 and 3, respectively. The top 20 pathways for each type are bolded in their respective type column. Pathways have been grouped to highlight the metabolic focus of each archetype.

| **Study** | **Subjects (n)** | **Samples (n)** | **Visits (n min)** | **Visits (n max)** | **Time span (min in days)** | **Time span (max in days)** |
| --- | --- | --- | --- | --- | --- | --- |
| PRJEB1220 | 11 | 22 | 2 | 2 | 55 | 76 |
| PRJEB1690 | 16 | 32 | 2 | 2 | 730 | 730 |
| PRJEB8094 | 3 | 7 | 2 | 3 | 7 | 90 |
| PRJNA290729 | 15 | 87 | 5 | 6 | 720 | 720 |
| PRJNA354235 | 163 | 425 | 2 | 3 | 2 | 180 |
| PRJNA491335 | 52 | 192 | 2 | 4 | 49 | 280 |
| WGM | 396 | 792 | 2 | 2 | 14 | 42 |

**Table S3.** Characteristics of studies with subjects having two or more visits, related to figure 5.

**References**

1.	Hale, V.L., Jeraldo, P., Mundy, M., Yao, J., Keeney, G., Scott, N., Cheek, E.H., Davidson, J., Greene, M., Martinez, C., et al. (2018). Synthesis of multi-omic data and community metabolic models reveals insights into the role of hydrogen sulfide in colon cancer. Methods 149, 59–68. https://doi.org/10.1016/j.ymeth.2018.04.024.

2.	Feng, Q., Liang, S., Jia, H., Stadlmayr, A., Tang, L., Lan, Z., Zhang, D., Xia, H., Xu, X., Jie, Z., et al. (2015). Gut microbiome development along the colorectal adenoma–carcinoma sequence. Nat. Commun. 6, 6528. https://doi.org/10.1038/ncomms7528.

3.	Yachida, S., Mizutani, S., Shiroma, H., Shiba, S., Nakajima, T., Sakamoto, T., Watanabe, H., Masuda, K., Nishimoto, Y., Kubo, M., et al. (2019). Metagenomic and metabolomic analyses reveal distinct stage-specific phenotypes of the gut microbiota in colorectal cancer. Nat. Med. 25, 968–976. https://doi.org/10.1038/s41591-019-0458-7.

4.	Zhang, X., Zhang, D., Jia, H., Feng, Q., Wang, D., Liang, D., Wu, X., Li, J., Tang, L., Li, Y., et al. (2015). The oral and gut microbiomes are perturbed in rheumatoid arthritis and partly normalized after treatment. Nat. Med. 21, 895–905. https://doi.org/10.1038/nm.3914.

5.	Qin, J., Li, Y., Cai, Z., Li, S., Zhu, J., Zhang, F., Liang, S., Zhang, W., Guan, Y., Shen, D., et al. (2012). A metagenome-wide association study of gut microbiota in type 2 diabetes. Nature 490, 55–60. https://doi.org/10.1038/nature11450.

6.	He, Q., Gao, Y., Jie, Z., Yu, X., Laursen, J.M., Xiao, L., Li, Y., Li, L., Zhang, F., Feng, Q., et al. (2017). Two distinct metacommunities characterize the gut microbiota in Crohn’s disease patients. GigaScience 6, gix050. https://doi.org/10.1093/gigascience/gix050.

7.	Fukuyama, J., Rumker, L., Sankaran, K., Jeganathan, P., Dethlefsen, L., Relman, D.A., and Holmes, S.P. (2017). Multidomain analyses of a longitudinal human microbiome intestinal cleanout perturbation experiment. PLoS Comput. Biol. 13, e1005706. https://doi.org/10.1371/journal.pcbi.1005706.

8.	Qin, J., Li, R., Raes, J., Arumugam, M., Burgdorf, K.S., Manichanh, C., Nielsen, T., Pons, N., Levenez, F., Yamada, T., et al. (2010). A human gut microbial gene catalog established by metagenomic sequencing. Nature 464, 59–65. https://doi.org/10.1038/nature08821.

9.	Karlsson, F.H., Tremaroli, V., Nookaew, I., Bergström, G., Behre, C.J., Fagerberg, B., Nielsen, J., and Bäckhed, F. (2013). Gut metagenome in European women with normal, impaired and diabetic glucose control. Nature 498, 99–103. https://doi.org/10.1038/nature12198.

10.	Jones, M.B., Highlander, S.K., Anderson, E.L., Li, W., Dayrit, M., Klitgord, N., Fabani, M.M., Seguritan, V., Green, J., Pride, D.T., et al. (2015). Library preparation methodology can influence genomic and functional predictions in human microbiome research. Proc. Natl. Acad. Sci. U. S. A. 112, 14024–14029. https://doi.org/10.1073/pnas.1519288112.

11.	Plichta, D.R., Somani, J., Pichaud, M., Wallace, Z.S., Fernandes, A.D., Perugino, C.A., Lähdesmäki, H., Stone, J.H., Vlamakis, H., Chung, D.C., et al. (2021). Congruent microbiome signatures in fibrosis-prone autoimmune diseases: IgG4-related disease and systemic sclerosis. Genome Med. 13, 35. https://doi.org/10.1186/s13073-021-00853-7.

12.	Raymond, F., Ouameur, A.A., Déraspe, M., Iqbal, N., Gingras, H., Dridi, B., Leprohon, P., Plante, P.-L., Giroux, R., Bérubé, È., et al. (2016). The initial state of the human gut microbiome determines its reshaping by antibiotics. ISME J. 10, 707–720. https://doi.org/10.1038/ismej.2015.148.

13.	Thomas, A.M., Manghi, P., Asnicar, F., Pasolli, E., Armanini, F., Zolfo, M., Beghini, F., Manara, S., Karcher, N., Pozzi, C., et al. (2019). Metagenomic analysis of colorectal cancer datasets identifies cross-cohort microbial diagnostic signatures and a link with choline degradation. Nat. Med. 25, 667–678. https://doi.org/10.1038/s41591-019-0405-7.

14.	King, C.H., Desai, H., Sylvetsky, A.C., LoTempio, J., Ayanyan, S., Carrie, J., Crandall, K.A., Fochtman, B.C., Gasparyan, L., Gulzar, N., et al. (2019). Baseline human gut microbiota profile in healthy people and standard reporting template. PLoS ONE 14, e0206484. https://doi.org/10.1371/journal.pone.0206484.

15.	Schirmer, M., Franzosa, E.A., Lloyd-Price, J., McIver, L.J., Schwager, R., Poon, T.W., Ananthakrishnan, A.N., Andrews, E., Barron, G., Lake, K., et al. (2018). Dynamics of metatranscription in the inflammatory bowel disease gut microbiome. Nat. Microbiol. 3, 337–346. https://doi.org/10.1038/s41564-017-0089-z.

16.	Guthrie, L., Gupta, S., Daily, J., and Kelly, L. (2017). Human microbiome signatures of differential colorectal cancer drug metabolism. Npj Biofilms Microbiomes 3, 1–8. https://doi.org/10.1038/s41522-017-0034-1.

17.	Yu, J., Feng, Q., Wong, S.H., Zhang, D., Liang, Q. yi, Qin, Y., Tang, L., Zhao, H., Stenvang, J., Li, Y., et al. (2017). Metagenomic analysis of faecal microbiome as a tool towards targeted non-invasive biomarkers for colorectal cancer. Gut 66, 70–78. https://doi.org/10.1136/gutjnl-2015-309800.

18.	Zeller, G., Tap, J., Voigt, A.Y., Sunagawa, S., Kultima, J.R., Costea, P.I., Amiot, A., Böhm, J., Brunetti, F., Habermann, N., et al. (2014). Potential of fecal microbiota for early-stage detection of colorectal cancer. Mol. Syst. Biol. 10, 766. https://doi.org/10.15252/msb.20145645.

19.	Pasolli, E., Asnicar, F., Manara, S., Zolfo, M., Karcher, N., Armanini, F., Beghini, F., Manghi, P., Tett, A., Ghensi, P., et al. (2019). Extensive Unexplored Human Microbiome Diversity Revealed by Over 150,000 Genomes from Metagenomes Spanning Age, Geography, and Lifestyle. Cell 176, 649-662.e20. https://doi.org/10.1016/j.cell.2019.01.001.

20.	Nielsen, H.B., Almeida, M., Juncker, A.S., Rasmussen, S., Li, J., Sunagawa, S., Plichta, D.R., Gautier, L., Pedersen, A.G., Le Chatelier, E., et al. (2014). Identification and assembly of genomes and genetic elements in complex metagenomic samples without using reference genomes. Nat. Biotechnol. 32, 822–828. https://doi.org/10.1038/nbt.2939.

21.	Rampelli, S., Schnorr, S.L., Consolandi, C., Turroni, S., Severgnini, M., Peano, C., Brigidi, P., Crittenden, A.N., Henry, A.G., and Candela, M. (2015). Metagenome Sequencing of the Hadza Hunter-Gatherer Gut Microbiota. Curr. Biol. 25, 1682–1693. https://doi.org/10.1016/j.cub.2015.04.055.

22.	Dhakan, D.B., Maji, A., Sharma, A.K., Saxena, R., Pulikkan, J., Grace, T., Gomez, A., Scaria, J., Amato, K.R., and Sharma, V.K. (2019). The unique composition of Indian gut microbiome, gene catalogue, and associated fecal metabolome deciphered using multi-omics approaches. GigaScience 8, giz004. https://doi.org/10.1093/gigascience/giz004.

23.	Wampach, L., Heintz-Buschart, A., Fritz, J.V., Ramiro-Garcia, J., Habier, J., Herold, M., Narayanasamy, S., Kaysen, A., Hogan, A.H., Bindl, L., et al. (2018). Birth mode is associated with earliest strain-conferred gut microbiome functions and immunostimulatory potential. Nat. Commun. 9, 5091. https://doi.org/10.1038/s41467-018-07631-x.

24.	Ye, Z., Zhang, N., Wu, C., Zhang, X., Wang, Q., Huang, X., Du, L., Cao, Q., Tang, J., Zhou, C., et al. (2018). A metagenomic study of the gut microbiome in Behcet’s disease. Microbiome 6, 135. https://doi.org/10.1186/s40168-018-0520-6.

25.	Vogtmann, E., Hua, X., Zeller, G., Sunagawa, S., Voigt, A.Y., Hercog, R., Goedert, J.J., Shi, J., Bork, P., and Sinha, R. (2016). Colorectal Cancer and the Human Gut Microbiome: Reproducibility with Whole-Genome Shotgun Sequencing. PLoS ONE 11, e0155362. https://doi.org/10.1371/journal.pone.0155362.

26.	Bedarf, J.R., Hildebrand, F., Coelho, L.P., Sunagawa, S., Bahram, M., Goeser, F., Bork, P., and Wüllner, U. (2017). Functional implications of microbial and viral gut metagenome changes in early stage L-DOPA-naïve Parkinson’s disease patients. Genome Med. 9, 39. https://doi.org/10.1186/s13073-017-0428-y.

27.	Qin, N., Yang, F., Li, A., Prifti, E., Chen, Y., Shao, L., Guo, J., Le Chatelier, E., Yao, J., Wu, L., et al. (2014). Alterations of the human gut microbiome in liver cirrhosis. Nature 513, 59–64. https://doi.org/10.1038/nature13568.

28.	Hansen, L.B.S., Roager, H.M., Søndertoft, N.B., Gøbel, R.J., Kristensen, M., Vallès-Colomer, M., Vieira-Silva, S., Ibrügger, S., Lind, M.V., Mærkedahl, R.B., et al. (2018). A low-gluten diet induces changes in the intestinal microbiome of healthy Danish adults. Nat. Commun. 9, 4630. https://doi.org/10.1038/s41467-018-07019-x.

29.	Molinaro, A., Bel Lassen, P., Henricsson, M., Wu, H., Adriouch, S., Belda, E., Chakaroun, R., Nielsen, T., Bergh, P.-O., Rouault, C., et al. (2020). Imidazole propionate is increased in diabetes and associated with dietary patterns and altered microbial ecology. Nat. Commun. 11, 5881. https://doi.org/10.1038/s41467-020-19589-w.

30.	Keohane, D.M., Ghosh, T.S., Jeffery, I.B., Molloy, M.G., O’Toole, P.W., and Shanahan, F. (2020). Microbiome and health implications for ethnic minorities after enforced lifestyle changes. Nat. Med. 26, 1089–1095. https://doi.org/10.1038/s41591-020-0963-8.

31.	Wirbel, J., Pyl, P.T., Kartal, E., Zych, K., Kashani, A., Milanese, A., Fleck, J.S., Voigt, A.Y., Palleja, A., Ponnudurai, R., et al. (2019). Meta-analysis of fecal metagenomes reveals global microbial signatures that are specific for colorectal cancer. Nat. Med. 25, 679–689. https://doi.org/10.1038/s41591-019-0406-6.

32.	Costea, P.I., Coelho, L.P., Sunagawa, S., Munch, R., Huerta‐Cepas, J., Forslund, K., Hildebrand, F., Kushugulova, A., Zeller, G., and Bork, P. (2017). Subspecies in the global human gut microbiome. Mol. Syst. Biol. 13, 960. https://doi.org/10.15252/msb.20177589.

33.	Tett, A., Huang, K.D., Asnicar, F., Fehlner-Peach, H., Pasolli, E., Karcher, N., Armanini, F., Manghi, P., Bonham, K., Zolfo, M., et al. (2019). The Prevotella copri Complex Comprises Four Distinct Clades Underrepresented in Westernized Populations. Cell Host Microbe 26, 666-679.e7. https://doi.org/10.1016/j.chom.2019.08.018.

34.	Mehta, R.S., Abu-Ali, G.S., Drew, D.A., Lloyd-Price, J., Subramanian, A., Lochhead, P., Joshi, A.D., Ivey, K.L., Khalili, H., Brown, G.T., et al. (2018). Stability of the human faecal microbiome in a cohort of adult men. Nat. Microbiol. 3, 347–355. https://doi.org/10.1038/s41564-017-0096-0.

35.	Rubel, M.A., Abbas, A., Taylor, L.J., Connell, A., Tanes, C., Bittinger, K., Ndze, V.N., Fonsah, J.Y., Ngwang, E., Essiane, A., et al. (2020). Lifestyle and the presence of helminths is associated with gut microbiome composition in Cameroonians. Genome Biol. 21, 122. https://doi.org/10.1186/s13059-020-02020-4.

36.	Yassour, M., Lim, M.Y., Yun, H.S., Tickle, T.L., Sung, J., Song, Y.-M., Lee, K., Franzosa, E.A., Morgan, X.C., Gevers, D., et al. (2016). Sub-clinical detection of gut microbial biomarkers of obesity and type 2 diabetes. Genome Med. 8, 17. https://doi.org/10.1186/s13073-016-0271-6.

37.	Le Chatelier, E., Nielsen, T., Qin, J., Prifti, E., Hildebrand, F., Falony, G., Almeida, M., Arumugam, M., Batto, J.-M., Kennedy, S., et al. (2013). Richness of human gut microbiome correlates with metabolic markers. Nature 500, 541–546. https://doi.org/10.1038/nature12506.

38.	Ianiro, G., Punčochář, M., Karcher, N., Porcari, S., Armanini, F., Asnicar, F., Beghini, F., Blanco-Míguez, A., Cumbo, F., Manghi, P., et al. (2022). Variability of strain engraftment and predictability of microbiome composition after fecal microbiota transplantation across different diseases. Nat. Med. 28, 1913–1923. https://doi.org/10.1038/s41591-022-01964-3.

39.	Hannigan, G.D., Duhaime, M.B., Ruffin, M.T., Koumpouras, C.C., and Schloss, P.D. (2018). Diagnostic Potential and Interactive Dynamics of the Colorectal Cancer Virome. mBio 9, 10.1128/mbio.02248-18. https://doi.org/10.1128/mbio.02248-18.

40.	Shao, Y., Forster, S.C., Tsaliki, E., Vervier, K., Strang, A., Simpson, N., Kumar, N., Stares, M.D., Rodger, A., Brocklehurst, P., et al. (2019). Stunted microbiota and opportunistic pathogen colonization in caesarean-section birth. Nature 574, 117–121. https://doi.org/10.1038/s41586-019-1560-1.

41.	David, L.A., Weil, A., Ryan, E.T., Calderwood, S.B., Harris, J.B., Chowdhury, F., Begum, Y., Qadri, F., LaRocque, R.C., and Turnbaugh, P.J. (2015). Gut Microbial Succession Follows Acute Secretory Diarrhea in Humans. mBio 6, e00381-15. https://doi.org/10.1128/mBio.00381-15.

42.	Louis, S., Tappu, R.-M., Damms-Machado, A., Huson, D.H., and Bischoff, S.C. (2016). Characterization of the Gut Microbial Community of Obese Patients Following a Weight-Loss Intervention Using Whole Metagenome Shotgun Sequencing. PLOS ONE 11, e0149564. https://doi.org/10.1371/journal.pone.0149564.

43.	Schirmer, M., Smeekens, S.P., Vlamakis, H., Jaeger, M., Oosting, M., Franzosa, E.A., ter Horst, R., Jansen, T., Jacobs, L., Bonder, M.J., et al. (2016). Linking the Human Gut Microbiome to Inflammatory Cytokine Production Capacity. Cell 167, 1125-1136.e8. https://doi.org/10.1016/j.cell.2016.10.020.

44.	Zhu, F., Ju, Y., Wang, W., Wang, Q., Guo, R., Ma, Q., Sun, Q., Fan, Y., Xie, Y., Yang, Z., et al. (2020). Metagenome-wide association of gut microbiome features for schizophrenia. Nat. Commun. 11, 1612. https://doi.org/10.1038/s41467-020-15457-9.

45.	Liu, W., Zhang, J., Wu, C., Cai, S., Huang, W., Chen, J., Xi, X., Liang, Z., Hou, Q., Zhou, B., et al. (2016). Unique Features of Ethnic Mongolian Gut Microbiome revealed by metagenomic analysis. Sci. Rep. 6, 34826. https://doi.org/10.1038/srep34826.

46.	Heintz-Buschart, A., May, P., Laczny, C.C., Lebrun, L.A., Bellora, C., Krishna, A., Wampach, L., Schneider, J.G., Hogan, A., de Beaufort, C., et al. (2016). Integrated multi-omics of the human gut microbiome in a case study of familial type 1 diabetes. Nat. Microbiol. 2, 1–13. https://doi.org/10.1038/nmicrobiol.2016.180.

47.	Xie, H., Guo, R., Zhong, H., Feng, Q., Lan, Z., Qin, B., Ward, K.J., Jackson, M.A., Xia, Y., Chen, X., et al. (2016). Shotgun Metagenomics of 250 Adult Twins Reveals Genetic and Environmental Impacts on the Gut Microbiome. Cell Syst. 3, 572-584.e3. https://doi.org/10.1016/j.cels.2016.10.004.

48.	Smits, S.A., Leach, J., Sonnenburg, E.D., Gonzalez, C.G., Lichtman, J.S., Reid, G., Knight, R., Manjurano, A., Changalucha, J., Elias, J.E., et al. (2017). Seasonal Cycling in the Gut Microbiome of the Hadza Hunter-Gatherers of Tanzania. Science 357, 802–806. https://doi.org/10.1126/science.aan4834.

49.	Nagy-Szakal, D., Williams, B.L., Mishra, N., Che, X., Lee, B., Bateman, L., Klimas, N.G., Komaroff, A.L., Levine, S., Montoya, J.G., et al. (2017). Fecal metagenomic profiles in subgroups of patients with myalgic encephalomyelitis/chronic fatigue syndrome. Microbiome 5, 44. https://doi.org/10.1186/s40168-017-0261-y.

50.	Ferretti, P., Pasolli, E., Tett, A., Asnicar, F., Gorfer, V., Fedi, S., Armanini, F., Truong, D.T., Manara, S., Zolfo, M., et al. (2018). Mother-to-Infant Microbial Transmission from Different Body Sites Shapes the Developing Infant Gut Microbiome. Cell Host Microbe 24, 133-145.e5. https://doi.org/10.1016/j.chom.2018.06.005.

51.	Bäckhed, F., Roswall, J., Peng, Y., Feng, Q., Jia, H., Kovatcheva-Datchary, P., Li, Y., Xia, Y., Xie, H., Zhong, H., et al. (2015). Dynamics and Stabilization of the Human Gut Microbiome during the First Year of Life. Cell Host Microbe 17, 690–703. https://doi.org/10.1016/j.chom.2015.04.004.

52.	Zeevi, D., Korem, T., Zmora, N., Israeli, D., Rothschild, D., Weinberger, A., Ben-Yacov, O., Lador, D., Avnit-Sagi, T., Lotan-Pompan, M., et al. (2015). Personalized Nutrition by Prediction of Glycemic Responses. Cell 163, 1079–1094. https://doi.org/10.1016/j.cell.2015.11.001.

53.	Bengtsson-Palme, J., Angelin, M., Huss, M., Kjellqvist, S., Kristiansson, E., Palmgren, H., Larsson, D.G.J., and Johansson, A. (2015). The Human Gut Microbiome as a Transporter of Antibiotic Resistance Genes between Continents. Antimicrob. Agents Chemother. 59, 6551–6560. https://doi.org/10.1128/AAC.00933-15.

54.	Sankaranarayanan, K., Ozga, A.T., Warinner, C., Tito, R.Y., Obregon-Tito, A.J., Xu, J., Gaffney, P.M., Jervis, L.L., Cox, D., Stephens, L., et al. (2015). Gut Microbiome Diversity among Cheyenne and Arapaho Individuals from Western Oklahoma. Curr. Biol. 25, 3161–3169. https://doi.org/10.1016/j.cub.2015.10.060.

55.	Li, J., Zhao, F., Wang, Y., Chen, J., Tao, J., Tian, G., Wu, S., Liu, W., Cui, Q., Geng, B., et al. (2017). Gut microbiota dysbiosis contributes to the development of hypertension. Microbiome 5, 14. https://doi.org/10.1186/s40168-016-0222-x.

56.	Jie, Z., Xia, H., Zhong, S.-L., Feng, Q., Li, S., Liang, S., Zhong, H., Liu, Z., Gao, Y., Zhao, H., et al. (2017). The gut microbiome in atherosclerotic cardiovascular disease. Nat. Commun. 8, 845. https://doi.org/10.1038/s41467-017-00900-1.

57.	Lokmer, A., Cian, A., Froment, A., Gantois, N., Viscogliosi, E., Chabé, M., and Ségurel, L. (2019). Use of shotgun metagenomics for the identification of protozoa in the gut microbiota of healthy individuals from worldwide populations with various industrialization levels. PLoS ONE 14, e0211139. https://doi.org/10.1371/journal.pone.0211139.

58.	Asnicar, F., Manara, S., Zolfo, M., Truong, D.T., Scholz, M., Armanini, F., Ferretti, P., Gorfer, V., Pedrotti, A., Tett, A., et al. (2017). Studying Vertical Microbiome Transmission from Mothers to Infants by Strain-Level Metagenomic Profiling. mSystems 2, e00164-16. https://doi.org/10.1128/mSystems.00164-16.

59.	Lloyd-Price, J., Mahurkar, A., Rahnavard, G., Crabtree, J., Orvis, J., Hall, A.B., Brady, A., Creasy, H.H., McCracken, C., Giglio, M.G., et al. (2017). Strains, functions and dynamics in the expanded Human Microbiome Project. Nature 550, 61–66. https://doi.org/10.1038/nature23889.

60.	Yassour, M., Jason, E., Hogstrom, L., Arthur, T.D., Tripathi, S., Siljander, H., Selvenius, J., Oikarinen, S., Hyöty, H., Virtanen, S.M., et al. (2018). Strain-level analysis of mother-to-child bacterial transmission during the first few months of life. Cell Host Microbe 24, 146-154.e4. https://doi.org/10.1016/j.chom.2018.06.007.

61.	Brito, I., Yilmaz, S., Huang, K., Xu, L., Jupiter, S., Jenkins, A., Naisilisili, W., Tamminen, M., Smillie, C., Wortman, J., et al. (2016). Mobile genes in the human microbiome are structured from global to individual scales. Nature 535, 435–439. https://doi.org/10.1038/nature18927.

62.	Ijaz, U.Z., Quince, C., Hanske, L., Loman, N., Calus, S.T., Bertz, M., Edwards, C.A., Gaya, D.R., Hansen, R., McGrogan, P., et al. (2017). The distinct features of microbial ‘dysbiosis’ of Crohn’s disease do not occur to the same extent in their unaffected, genetically-linked kindred. PLoS ONE 12, e0172605. https://doi.org/10.1371/journal.pone.0172605.

63.	Fromentin, S., Forslund, S.K., Chechi, K., Aron-Wisnewsky, J., Chakaroun, R., Nielsen, T., Tremaroli, V., Ji, B., Prifti, E., Myridakis, A., et al. (2022). Microbiome and metabolome features of the cardiometabolic disease spectrum. Nat. Med. 28, 303–314. https://doi.org/10.1038/s41591-022-01688-4.

64.	Maldonado-Contreras, A., Ferrer, L., Cawley, C., Crain, S., Bhattarai, S., Toscano, J., Ward, D.V., and Hoffman, A. (2020). Dysbiosis in a canine model of human fistulizing Crohn’s disease. Gut Microbes 12, 1785246. https://doi.org/10.1080/19490976.2020.1785246.

65.	Li, S.S., Zhu, A., Benes, V., Costea, P.I., Hercog, R., Hildebrand, F., Huerta-Cepas, J., Nieuwdorp, M., Salojärvi, J., Voigt, A.Y., et al. (2016). Durable coexistence of donor and recipient strains after fecal microbiota transplantation. Science 352, 586–589. https://doi.org/10.1126/science.aad8852.

66.	De Angelis, M., Ferrocino, I., Calabrese, F.M., De Filippis, F., Cavallo, N., Siragusa, S., Rampelli, S., Di Cagno, R., Rantsiou, K., Vannini, L., et al. (2020). Diet influences the functions of the human intestinal microbiome. Sci. Rep. 10, 4247. https://doi.org/10.1038/s41598-020-61192-y.

67.	Hall, A.B., Yassour, M., Sauk, J., Garner, A., Jiang, X., Arthur, T., Lagoudas, G.K., Vatanen, T., Fornelos, N., Wilson, R., et al. (2017). A novel Ruminococcus gnavus clade enriched in inflammatory bowel disease patients. Genome Med. 9, 103. https://doi.org/10.1186/s13073-017-0490-5.

68.	Rosa, B.A., Supali, T., Gankpala, L., Djuardi, Y., Sartono, E., Zhou, Y., Fischer, K., Martin, J., Tyagi, R., Bolay, F.K., et al. (2018). Differential human gut microbiome assemblages during soil-transmitted helminth infections in Indonesia and Liberia. Microbiome 6, 33. https://doi.org/10.1186/s40168-018-0416-5.

69.	Vincent, C., Miller, M.A., Edens, T.J., Mehrotra, S., Dewar, K., and Manges, A.R. (2016). Bloom and bust: intestinal microbiota dynamics in response to hospital exposures and Clostridium difficile colonization or infection. Microbiome 4, 12. https://doi.org/10.1186/s40168-016-0156-3.

70.	De Filippis, F., Pasolli, E., Tett, A., Tarallo, S., Naccarati, A., Angelis, M.D., Neviani, E., Cocolin, L., Gobbetti, M., Segata, N., et al. (2019). Distinct Genetic and Functional Traits of Human Intestinal Prevotella copri Strains Are Associated with Different Habitual Diets. Cell Host Microbe 25, 444-453.e3. https://doi.org/10.1016/j.chom.2019.01.004.
